# Unsupervised protein embeddings outperform hand-crafted sequence and structure features at predicting molecular function

**DOI:** 10.1101/2020.04.07.028373

**Authors:** Amelia Villegas-Morcillo, Stavros Makrodimitris, Roeland C.H.J. van Ham, Angel M. Gomez, Victoria Sanchez, Marcel J.T. Reinders

**Affiliations:** Dept. of Signal Theory, Telematics and Communications, University of Granada, Calle Periodista Daniel Saucedo Aranda, 18071, Granada, Spain; Delft Bioinformatics Lab, Delft University of Technology, Van Mourik Broekmanweg 6, 2628XE, Delft, the Netherlands; Keygene N.V., Agro Business Park 90, 6708PW, Wageningen, the Netherlands; Leiden Computational Biology Center, Leiden University Medical Center, Einthovenweg 20, 2333ZC, Leiden, the Netherlands

**Author notes:** Equal contribution (Amelia Villegas-Morcillo, Stavros Makrodimitris).

## Abstract

**Motivation:** Protein function prediction is a difficult bioinformatics problem. Many recent methods use deep neural networks to learn complex sequence representations and predict function from these. Deep supervised models require a lot of labeled training data which are not available for this task. However, a very large amount of protein sequences without functional labels is available.

**Results:** We applied an existing deep sequence model that had been pre-trained in an unsupervised setting on the supervised task of protein function prediction. We found that this complex feature representation is effective for this task, outperforming hand-crafted features such as one-hot encoding of amino acids, *k*-mer counts, secondary structure and backbone angles. Also, it partly negates the need for deep prediction models, as a two-layer perceptron was enough to achieve state-of-the-art performance in the third Critical Assessment of Functional Annotation benchmark. We also show that combining this sequence representation with protein 3D structure information does not lead to performance improvement, hinting that three-dimensional structure is also potentially learned during the unsupervised pre-training.

**Availability:** Implementations of all used models can be found at https://github.com/stamakro/GCN-for-Structure-and-Function.

**Supplementary information:** Supplementary data are available online.

## 1 Introduction

Proteins perform most of the functions necessary for life. However, proteins with a well-characterized function are only a small fraction of all known proteins and mostly restricted to a few model species. Therefore, the ability to accurately predict protein function has the potential to accelerate research in fields such as animal and plant breeding, biotechnology, and human health.

The most common data type used for automated function prediction (AFP) is the amino acid sequence, as conserved sequence implies conserved function (Kimura and Ohta, 1974). Consequently, many widely-used AFP algorithms rely on sequence similarity search via BLAST (Altschul *et al*., 1990) and its variants or on hidden Markov models (Eddy, 2009). Other types of sequence information that have been used include *k*-mer counts, predicted secondary structure, sequence motifs, conjoint triad features and pseudo-amino acid composition (Cozzetto *et al*., 2016; Fa *et al*., 2018; Sureyya Rifaioglu *et al*., 2019). Moreover, Cozzetto et al. showed that different sequence features are informative for different functions.

More recently, advances in machine learning have partially shifted the focus from hand-crafted features, such as those described above, to automatic representation learning, where a complex model-most often a neural network-is used to learn features that are useful for the prediction task at hand. Many such neural network methods have been proposed, which use a variety of architectures (Bonetta and Valentino, 2019).

Some studies combined the two approaches, starting from hand-crafted features that are fed into a multi-layer perceptron (MLP) to learn more elaborate representations (Fa *et al*., 2018; Sureyya Rifaioglu *et al*., 2019). Others apply recurrent or convolutional architectures to directly process variable-length sequences. For instance, (Kulmanov *et al*., 2018) used a neural embedding layer to embed all possible amino acid triplets into a 128-dimensional space and then applied a convolutional neural network (CNN) on these triplet embeddings. Moreover, (Liu, 2017) and (Cao *et al*., 2017) trained Long Short-Term Memory (LSTM) networks to perform AFP.

The motivation behind these deep models is that functional information is encoded in the sequence in a complicated way. A disadvantage is that complex models with a large number of parameters require a large amount of training examples, which are not available for the AFP task. There are about 80,000 proteins with at least one experimentally-derived Molecular Function Gene Ontology (GO) (Ashburner *et al*., 2000) annotation in SwissProt and 11,123 terms in total.

On the other hand, a huge number of protein sequences of unknown function is available (>175M in UniProtKB). Although these sequences cannot be directly used to train an AFP model, they can be fed into an unsupervised deep model that tries to learn general amino acid and/or protein features. This learned representation can then be applied to other protein-related tasks, including AFP, either directly or after fine-tuning by means of supervised training. Several examples of unsupervised pre-training leading to substantial performance improvement exist in the fields of computer vision (Doersch *et al*., 2015; Gidaris *et al*., 2018; Mathis *et al*., 2019) and natural language processing (NLP) (McCann *et al*., 2017; Peters *et al*., 2018; Devlin *et al*., 2018).

A deep unsupervised model of protein sequences was recently made available (Heinzinger *et al*., 2019). It is based on the NLP model ELMo (Embeddings from Language Models) (Peters *et al*., 2018) and is composed of a character-level CNN (CharCNN) followed by two layers of bidirectional LSTMs. The CNN embeds each amino acid into a latent space, while the LSTMs use that embedding to model the context of the surrounding amino acids. The hidden states of the two LSTM layers and the latent representation are added to give the final context-aware embedding. These embeddings demonstrated competitive performance in both amino acid and protein classification tasks, such as inferring the protein secondary structure, structural class, disordered regions, and cellular localization (Heinzinger *et al*., 2019; Kane *et al*., 2019). Other works also trained LSTMs to predict the next amino acid in a protein sequence using the LSTM hidden state at each amino acid as a feature vector (Gligorijevic *et al*., 2019; Alley *et al*., 2019). Finally, a transformer neural network was trained on 250 million protein sequences, yielding embeddings that reflected both protein structure and function (Rives *et al*., 2019).

Protein function is encoded in the amino acid sequence, but sequences can diverge during evolution while maintaining the same function. Protein structure is also known to determine function and is-in principle-more conserved than sequence (Wilson *et al*., 2000; Weinhold *et al*., 2008). From an AFP viewpoint, two proteins with different sequences can be assigned with high confidence to the same function if their structures are similar. It is therefore generally thought that combining sequence data with 3D structure leads to more accurate function predictions for proteins with known structure, especially for those without close homologues.

Structural information is often encoded as a protein distance map. This is a symmetric matrix containing the Euclidean distances between pairs of residues within a protein and is invariant to translations or rotations of the molecule in 3D space. One can obtain a binary representation from this real-valued matrix, called protein contact map, by applying a distance threshold (typically from 5 to 20 Å). This two-dimensional representation successfully captures the overall protein structure (Bartoli *et al*., 2007; Duarte *et al*., 2010). The protein contact map can be viewed as a binary image, where each pixel indicates whether a specific pair of residues are in contact or not. Alternatively, it can be interpreted as the adjacency matrix of a graph, where each amino acid is a node and edges represent amino acids that are in contact with each other. In order to extract meaningful information from contact maps, both two-dimensional CNNs (Zhu *et al*., 2017; Zheng *et al*., 2019) and graph convolutional networks (GCNs) (Fout *et al*., 2017; Zamora-Resendiz and Crivelli, 2019) have been proposed.

Only (Gligorijevic *et al*., 2019) have explored the effectiveness of a pre-trained sequence model in AFP, but it was done in combination with protein structure information using a GCN. We suspect that a deep pre-trained embedding can be powerful enough to predict protein function, in which case the structural information would not offer any significant performance improvement. Therefore, we set out to evaluate pre-trained ELMo embeddings in the task of predicting molecular functions, by comparing them to hand-crafted sequence and structural features in combination with 3D structure information in various forms. Fig. 1 provides an overview of the data and models used in our experiments. We demonstrate the effectiveness of the ELMo model (Heinzinger *et al*., 2019) and show that protein structure does not provide a significant performance boost to these embeddings, although it does so when we only consider a simple protein representation based on one-hot encoded amino acids.

**Fig. 1.**
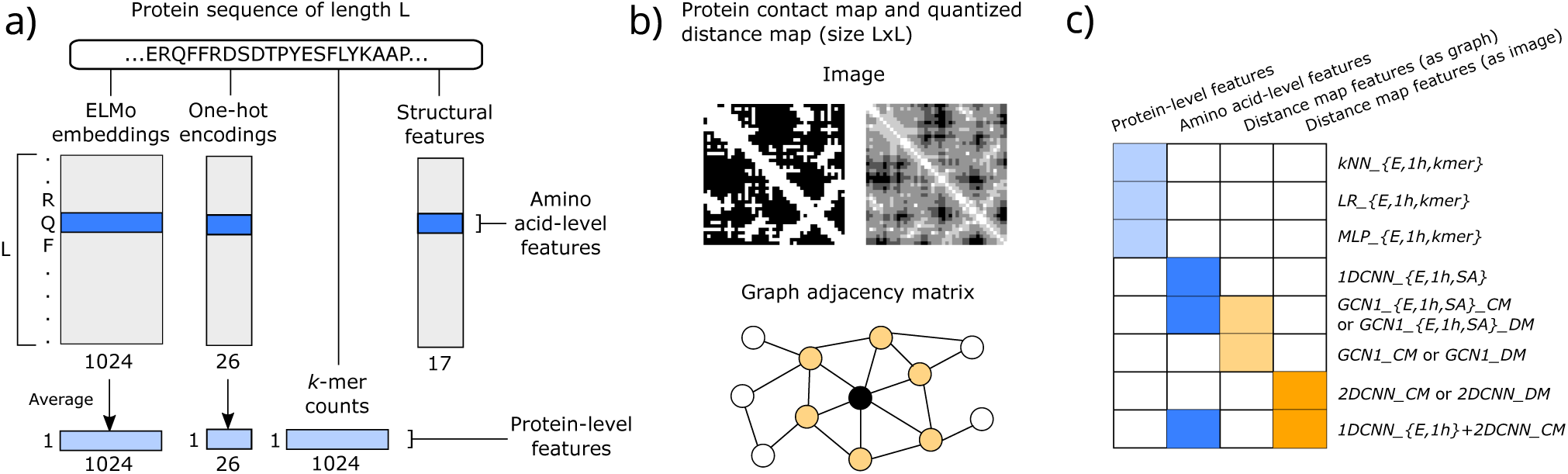
Protein representation types considered in this study, which encode (a) amino acid sequence information (ELMo embeddings, one-hot encodings, k-mer counts, and secondary structure and backbone angles) and (b) 3D structure information in the form of contact/distance map (as an image or graph adjacency matrix). (c) The protein representations (columns) that are fed as input to each classification model (rows) are indicated by a shaded box, colored blue for sequence and orange for distance map representations.

## 2 Materials & Methods

### 2.1 Protein representations

We considered two types of representations of the proteins (Fig. 1). The first one describes the sequence using amino acid features and the second one the three-dimensional structure, mainly in the form of contact maps.

For each sequence of length L, we extracted *amino acid-level features* using a pre-trained unsupervised language model (Heinzinger *et al*., 2019). This model is based on ELMo (Peters *et al*., 2018) and outputs a feature vector of dimension d=1,024 for each amino acid in the sequence. We denote this as a matrix X^*E*^ ∈ ℝ^*L*×*d*^. As proposed in (Heinzinger *et al*., 2019), we also obtained a fixed-length vector representation of each protein (*protein-level features*, denoted as x^*E*^ ∈ ℝ^*d*^) by averaging each feature over the L amino acids.

To compare ELMo with simpler sequence representations, we used the one-hot encoding of the amino acids, denoted by the matrix X^1*h*^ ∈ {0, 1}^*L*×*d*^ with d=26. As before, we obtained a protein-level representation x^1*h*^ ∈ R^*d*^, which contains the frequency of each amino acid in the protein sequence, completely ignoring the order. We also used a protein-level representation based on *k*-mer counts. To reduce the dimensionality of this representation, we applied truncated singular value decomposition (SVD) keeping the first 1,024 components (x^*kmer*^ ∈ R^*d*^).

With respect to structural information, we considered the protein distance map. This L×L matrix contains the Euclidean distances between all pairs of beta carbon atoms (alpha carbon atoms for Glycine) within each protein chain. We converted this matrix to a binary contact map using a threshold of 10 Å. We also tested an amino acid-level structural representation X^*SA*^ ∈ R^*L*×*d*^, with d=17 features including the secondary structure state and backbone angles. More details about the protein representations, as well as alternative ways of thresholding the distance maps can be found in Supplementary Material 1 (SM1).

### 2.2 Function prediction methods

In our experiments, we trained and evaluated several classifiers which use the protein representations defined above (Fig. 1). Details concerning hyperparameters and training are provided in SM2 (Tables S1-2).

We first considered methods operating on the protein-level features (either ELMo embeddings x^*E*^, one-hot encodings x^1*h*^, or *k*-mer counts x^*kmer*^). As these feature vectors are of fixed size for all proteins, we can apply traditional machine learning algorithms. Here, we tested the following classifiers: *k*-nearest neighbors (*k*-NN) with Euclidean distance, logistic regression (LR) with L2 regularization, and multi-layer perceptron (MLP) with one hidden layer. We denoted these models as *kNN_{E,1h,kmer}, LR_{E,1h,kmer}* and *MLP_{E,1h,kmer}*, respectively.

We also trained several convolutional networks on the amino acid-level representations (X^*E*^, X^1*h*^ or structural features X^*SA*^) and distance map data. The architectures are composed of convolutional layers; either 1D, 2D or graph-based. As the input size is variable in the sequence dimension, these layers are followed by a global pooling operation, to obtain a fixed-size vector for each protein. This embedding vector is then used to predict the corresponding C outputs (GO terms) through fully-connected (FC) layers. In the output layer we applied the sigmoid function, so that the final prediction for each GO term is in the range [0,1]. We tested either one or two FC layers and selected the optimal for each model based on the validation set. As shown in Table S3 (SM2), the architecture with one FC layer was preferred for all networks in the presence of ELMo embeddings, and the one with two FC layers when using one-hot encodings or contact map information only.

The one-dimensional convolutional neural network (1D-CNN) applies dilated convolutions in two layers (Fig. S1) and we refer to this model as *1DCNN_{E,1h,SA}*. To incorporate contact map information, we trained GCN models. In this case the protein 3D structure is viewed as a graph with adjacency matrix A ∈ {0, 1}^*L*×*L*^, where each amino acid of the sequence corresponds to a node and an edge between two nodes denotes that they are in contact. The graph convolution operator that we mainly used was the first-order approximation of the spectral graph convolution defined in (Kipf and Welling, 2019) as:

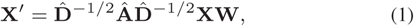

where Â = A + I is the adjacency matrix with self-loops, 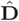 the diagonal degree matrix with 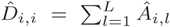 and W the weight matrix that combines the node features. Equation (1) describes the diffusion of information about each amino acid to the neighboring residues, where the neighborhood is defined by the graph. We tested the model proposed by (Gligorijevic *et al*., 2019) that has three convolutional layers (*GCN3_{E,1h,SA}_CM*, Fig. S2). As we intended to use simple models, we also considered a reduced version of this network, with only one convolutional layer (*GCN1_{E,1h,SA}_CM*, Fig. S3). In addition, GCN models can easily handle distance maps, by making use of weighted edges in the graph. Therefore, we trained both GCN models replacing the contact maps with quantized distance maps (*GCN3_{E,1h}_DM* and *GCN1_{E,1h}_DM*). The optimal number of bins in the quantized map was selected based on the validation set, as shown in Table S4 (SM2). We also tested three other graph convolution operators, which are described in SM3.

In order to test the ability of predicting function based on distance/contact maps alone, we evaluated two alternative approaches. The first one is based on the GCN model described above (Kipf and Welling, 2019) keeping A as before, but with X ∈ R^*L*×1^ containing the degree of each node as amino acid feature. Therefore, by applying the convolution operation of equation (1), the network only learns graph connectivity patterns (*GCN1_CM* and *GCN1_DM*). The second approach processes the maps as L × L images and learns image patterns using a 2D-CNN model with two convolutional layers (Fig. S4). We denoted this model as *2DCNN_CM* or *2DCNN_DM*, if the input is the contact map or the distance map, respectively.

Moreover, we investigated alternative ways of combining sequence and structure information, such as a combined 1D-CNN and 2D-CNN model that is simultaneously trained to extract a joint representation (Fig. S5). In this case, we concatenated the outputs of the two convolutional parts before the global pooling layer. We refer to this model as *1DCNN_E+2DCNN_CM* and *1DCNN_1h+2DCNN_CM*.

Finally, as baseline methods we used the naive (Radivojac *et al*., 2013) and BLAST (Altschul *et al*., 1990) methods. The naive method assigns a GO term to all test proteins with a probability equal to the frequency of that term in the training set. BLAST annotates each protein with the GO annotations of its top BLAST hit.

### 2.3 Data

We compared models that only use sequence information to models that also include contact maps. To do so, we considered proteins whose structure is available in the Protein Data Bank (Berman *et al*., 2000). We refer to this dataset as *PDB*. To better assess the sequence-only models, we also applied them to a larger dataset (referred to as *SP*) that includes all proteins from the SwissProt database. Finally, we also evaluated the ELMo models on the CAFA3 benchmark (Zhou *et al*., 2019) (*CAFA* dataset).

For the *PDB* dataset, we retrieved all protein chains with known PDB structure and for *SP* all sequences that were available in SwissProt in January 2020. We only considered proteins with sequence length in the range [40, 1000] that had GO annotations in the Molecular Function Ontology (MFO) with non-computational evidence codes. We used CD-HIT (Fu *et al*., 2012) to remove redundant sequences with an identity threshold of 95%. After these filtering steps, we had a total of 11,749 protein chains in *PDB* and 80,176 protein sequences in the *SP* dataset. To ensure diversity in the evaluation, we clustered all the protein sequences in each dataset with PSI-CD-HIT (Fu *et al*., 2012) into groups of maximum 30% identity, from which we randomly selected the samples to use as test set (10% of the entire dataset). The remaining data were randomly split into a training (80% of total proteins) and a validation set (10% of total proteins). We further defined a subset of the test set using BLAST, in which all proteins had sequence identity smaller than 30% to any of the training proteins. We excluded GO terms that had fewer than 40 positive examples in the training set or fewer than 5 in the validation or test sets and removed proteins that had no annotations after this filtering. Finally, our training, validation and test sets for *PDB* had 9,395, 1,173 and 450 proteins, respectively, annotated with C=256 MFO GO terms. In the *SP* dataset, we had 63,994 training, 8,004 validation and 3,530 test proteins, annotated with C=441 terms. More details about the data pre-processing steps are provided in SM4.

The *CAFA* training and test sets were provided by the organizers (Zhou *et al*., 2019). The test set contains 454 proteins. We randomly split the given training set into 90% for training (28,286 proteins) and 10% for validation (3,143 proteins), annotated with C=679 MFO GO terms. We did not apply sequence similarity filters on the *CAFA* dataset, as in that case we intend to exploit information present in closely related proteins.

### 2.4 Performance evaluation

The performance was measured using the maximum protein-centric F-measure (F_*max*_), the normalized minimum semantic distance (S_*min*_) (Clark and Radivojac, 2013; Jiang *et al*., 2016) and the term-centric *ROCAUC*. We estimated 95% confidence intervals (CI’s) using bootstrapping: we drew random samples with replacement from the test set until we obtained a set of proteins with a size equal to the original test set and calculated the metric values in this new set. We repeated this procedure 1,000 and 100 times for the *PDB* and *SP* test sets respectively. We used the same bootstrap sets for all methods, enabling joint comparisons across the bootstraps.

## 3 Results

### 3.1 Deep, pre-trained embeddings outperform hand-crafted sequence representations

We first compared the unsupervised ELMo embeddings of protein sequences to hand-crafted sequence representations at the task of predicting MFO terms, using the 30% sequence identity *PDB* test subset (450 protein chains). We used the amino acid-level features (X^*E*^, X^1*h*^ and X^*SA*^) in a 1D-CNN model and two GCN models, and compared to the *k*-NN, logistic regression (LR) and multi-layer perceptron (MLP) classifiers, which use the protein-level features (x^*E*^, x^1*h*^ and x^*kmer*^). As seen in Fig. 2 and Table S6 (SM5), the models using ELMo embeddings significantly outperform their counterparts using other features on all evaluation metrics. One exception is *GCN1_1h_CM*, which performs similarly to *GCN1_E_CM* in terms of *ROCAUC*. It achieves the top performance among models that use X^1*h*^, for all three metrics, although *MLP_1h* achieved equal F*max* (Table S6, SM5). Also, the protein-level x^1*h*^ representation (*k*-mers with k=1) consistently outperformed x^*kmer*^ which uses larger k values.

**Fig. 2.**
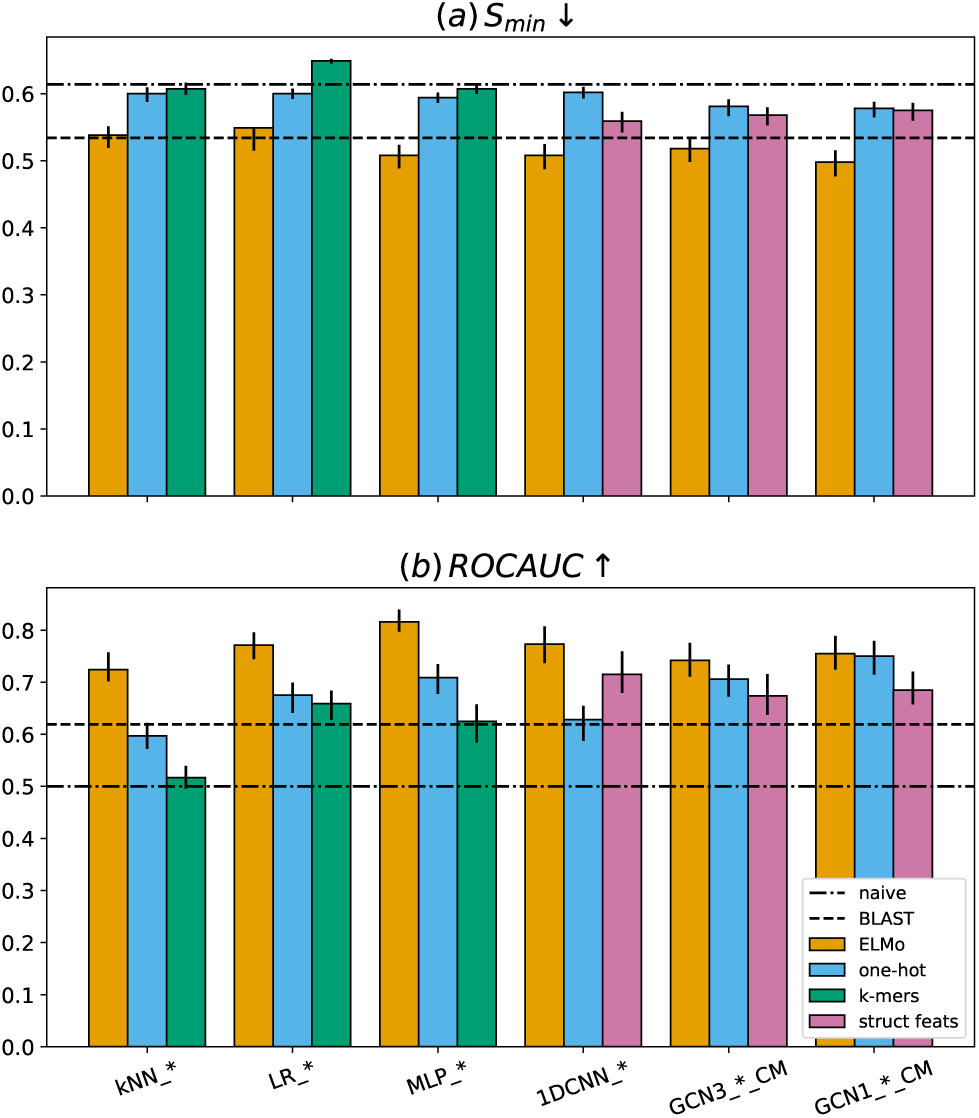
*S*_*min*_ (a) and *ROCAUC* (b) of models trained using either ELMo embeddings (orange), one-hot encodings (blue), k-mer counts (green) or structural features (pink), on the 30% sequence identity *PDB* test subset. The arrows denote that lower values (in *S*_*min*_) and higher values (in *ROCAUC*) correspond to better performance. The error bars denote 95% confidence intervals estimated using 1,000 bootstraps. The dashed line corresponds to the performance of BLAST and the dashed dotted line to the naive baseline.

We also evaluated the sequence-only models in the larger *SP* dataset (3,530 test proteins, 441 terms) and observed a similar pattern (Table S7, SM6). The absolute performances are better, but the superiority of ELMo embeddings is evident, as even simple models such as *kNN_E* and *LR_E* outperform all more complex models that use one-hot encodings. Analyzing the performance per GO term, we found that although *kNN_E* has a larger mean *ROCAUC* than *1DCNN_1h*, its superiority is mainly shown on the most frequent terms (Figs. S6a-b, SM7). On the contrary, all other tested models that use ELMo embeddings tend to have better performance for more specific terms (Fig. S7, SM7) and they consistently outperform the one-hot encodings-based models across all levels of the GO graph (Figs. S6c-h, SM7).

To get an additional evaluation of the ELMo embeddings compared to the state-of-the-art, we used them in the *CAFA* dataset (454 test proteins, 679 terms). Table S8 (SM6) shows the performance of *kNN_E, LR_E, MLP_E* and *1DCNN_E* in this dataset. All had quite competitive performance, outperforming at least 80% of the methods participating in CAFA3 (Zhou *et al*., 2019), while having 100% coverage, meaning that they could make predictions for all test proteins. Our top model, *MLP_E*, achieved an F*max* of 0.55, outperforming all but 4 of the methods that had participated in the challenge (Table S8, SM6).

### 3.2 GCN performs similarly to CNN when using ELMo embeddings

We then tested whether combining the ELMo embeddings with contact map information in a GCN improves the performance, for which we considered the *PDB* dataset. Fig. 2 and Table S6 (SM5) show the results in terms of F_*max*_, normalized S_*min*_ and *ROCAUC*, while Fig. S8 (SM8) shows the variance of the performance values estimated using bootstraps. Figs. S9-S11 (SM8) show all pairwise comparisons between the tested methods. The 3-layer GCN proposed in (Gligorijevic *et al*., 2019) trained with the ELMo embeddings (*GCN3_E_CM*) was worse than the *1DCNN_E* based on all three metrics (S_*min*_=0.52, *ROCAUC*=0.75, F_*max*_=0.48, compared to S_*min*_=0.51, *ROCAUC*=0.77, F_*max*_=0.50).

We also tested whether a simpler GCN model would be more efficient. We found that just a one layer graph convolutional network (*GCN1_E_CM*) outperformed the deeper GCN model on all metrics (Fig. 2 and Table S6, SM5). In fact, *GCN1_E_CM* had slightly better S_*min*_ than the *1DCNN_E* (2% improvement), though worse *ROCAUC*.

Furthermore, we compared these models that use amino acid-level ELMo embeddings to standard classifiers that use the protein-level embeddings. We observed that *LR_E* achieved similar *ROCAUC* to the *1DCNN_E* (0.77). *LR_E* achieved a S_*min*_ of 0.55, while *1DCNN_E* was about 7% better with 0.51. Despite the small difference, *1DCNN_E* outperformed *LR_E* in 998 out of 1,000 the bootstraps denoting that the difference is significant (Fig. S10b, SM8). Note that the test S_*min*_ of *LR_E* was worse than the performance achieved by the same model in the vast majority of the bootstraps (error bar in Fig. 2). The *kNN_E* had comparable S_*min*_ to *LR_E* and worse *ROCAUC* than all. The two-layer MLP on the protein-level embeddings (*MLP_E*) achieved the best *ROCAUC*, yielding a 5.3% improvement upon the second best method (*1DCNN_E*). These two models had equal S_*min*_ (Fig. 2 and Fig. S10e, SM8). Finally, *MLP_E* and *GCN1_E_CM* had the best F*max* (0.51), followed closely by *1DCNN_E* (0.50).

Comparing all the models jointly, *GCN1_E_CM* had the best S_*min*_ in 84.2% of the bootstraps followed by *1DCNN_E* with 9.5% and *MLP_E* with 6.2%. So, if we were able to sample future test sets from the same distribution, we expect *GCN1_E_CM* to have the smallest S_*min*_ 84.2% of the time. The conclusion is different when we evaluate on the term-centric *ROCAUC*, with *MLP_E* being the best in all bootstraps. These mixed results of the GCN and 1D-CNN, and the small differences to simple models on the protein-level embeddings hint that the performance of all tested models mainly stems from the power of the ELMo embeddings and not from the convolutions. To ensure that our observation about GCNs does not depend on the choice of the graph convolution operator, we repeated the experiments using other three graph operators and obtained similar results (Table S5, SM3).

### 3.3 Protein structure does not add information to the ELMo embeddings

In order to explain the lack of significant improvement when including the contact map information, we investigated the behavior of the GCN further, focusing on the 1-layer model, which was the better of the two tested GCNs. Keeping the architecture the same, we re-trained and tested the model, replacing each contact map with (a) a disconnected graph, i.e. substituting A with the identity matrix (*GCN1_E_I*), and (b) a random undirected graph with the same number of edges as the original (*GCN1_E_R*). As shown in Table S6 (SM5), the performance remains the same as that of the original contact map for both perturbations of the graphs, hinting that the sequence embeddings are enough for learning a good functional representation. However, replacing ELMo with one-hot encodings in this experiment (*GCN1_1h_I* and *GCN1_1h_R*) led to a performance drop compared to *GCN1_1h_CM* (Table S6, SM5).

We then trained a GCN model without using sequence features (*GCN1_CM*), “forcing” the network to learn to differentiate among the different GO terms using only the contact map. The results in Table 1 and Table S6 (SM5) show that the performance of that network was remarkably worse than *GCN1_1h_CM*, having an S_*min*_ of 0.60 and *ROCAUC* of 0.64. To put these numbers into perspective, the simple BLAST baseline had S_*min*_ of 0.53 and *ROCAUC* of 0.62. On the other hand, modeling the contact maps as images and not as graphs and feeding them into a custom 2D-CNN (*2DCNN_CM*) achieved better performance (S_*min*_ = 0.58 and *ROCAUC* = 0.68), although significantly worse than the models that used sequence features. Furthermore, the combined *1DCNN_E+2DCNN_CM* did not outperform *1DCNN_E*, and *1DCNN_1h+2DCNN_CM* was worse than *2DCNN_CM* (Table S6, SM5), showing that integrating sequence and structural features is not trivial.

**Table 1.**
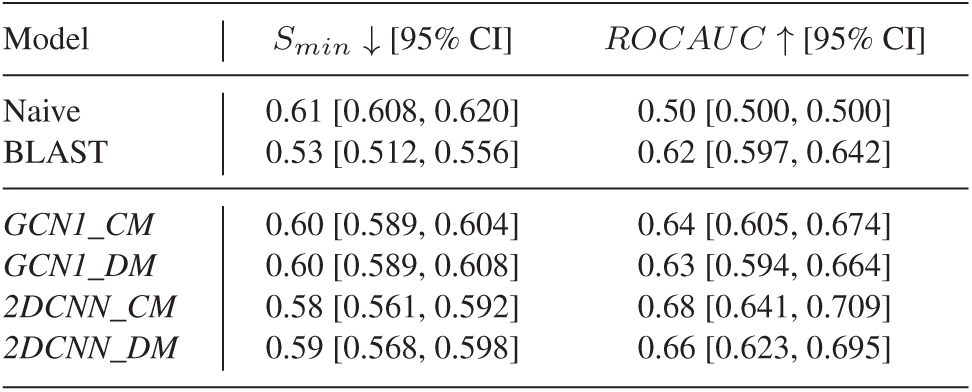
*S*_*min*_ and *ROCAUC* of the GCN and 2D-CNN networks that only use distance map information, compared to the naive and BLAST classifiers. All networks were evaluated using the 30% sequence identity *PDB* test subset. The 95% confidence intervals were estimated using 1,000 bootstraps

We then replaced the contact map with a quantized distance map, which led to mostly equal or worse performance (Table 1 and Table S6, SM5). All these results demonstrate that although contact and distance maps can in general be used for AFP, in the presence of ELMo embeddings they are not particularly useful. In another attempt to represent structure differently, we trained the 1D-CNN model with amino acid-level structural features based on secondary structure and backbone angles (X^*SA*^). They performed better than the one-hot encodings, but considerably worse than the ELMo embeddings (Fig. 2 and Table S6, SM5).

### 3.4 Supervised protein embeddings give insights into the behavior of the models

To better understand the differences between the models, we compared the embeddings learned by each of them. We fed all trained models with every protein from our *PDB* dataset and saved the 512-dimensional embedding vector after the global pooling layer, which gave us an 11, 740 × 512 embedding matrix. We then calculated the rank of each of these matrices to assess how “rich” the learned representations are. As shown in Table S9 (SM9), all methods that use the ELMo representation are either full-rank or very close to full-rank (508-512). On the other hand, the models that only operated on contact maps learned much simpler, lower-dimensional representations, with rank 310 for *2DCNN_CM* and 105 for *GCN1_CM*. By applying principal components analysis (PCA) to the *GCN1_CM* embeddings, we found that out of the 3 components explained 99.8% of the total variance (Fig. S12, SM9), suggesting that essentially this network learned a 3-feature representation of the proteins.

We also compared the embeddings of the different supervised models to the unsupervised ELMo embeddings. For every pair of test-training proteins from our *PDB* dataset, we calculated their cosine similarity in the embedding space, as well as a measure of similarity of their GO annotations based on the Jaccard index (Pesquita *et al*., 2007). For the ELMo embeddings, we found that the two similarity measures were significantly correlated (Fig. S13, with Spearman ρ=0.07, permutation p-value < 10^−4^, SM10). By extracting the embeddings from a supervised model such as *1DCNN_E* and *MLP_E*, the correlation value doubled (ρ=0.14, p-value < 10^−4^, SM10). For the *GCN1_E_CM*, the correlation value was 0.11 (Fig. S14, SM10). This verifies that unsupervised pre-training is able to capture some information about protein function, while additional supervised training provides extra information to the model.

Finally, to test to what extent different models learn similar embeddings, we clustered them based on the overlap of their 40 nearest neighborhood graphs, measured using Jaccard distance (Fig. 3 and SM11). We observed that the embeddings of *MLP_E* are the most similar to ELMo (Jaccard distance of 0.68, meaning that about one third of the 40 nearest neighbors are common). The models that used a 1-layer GCN (*GCN1_E_CM, GCN1_E_I* and *GCN1_E_R*) learned relatively similar neighborhoods to each other, clustering together at distance 0.77. Moreover, all ELMo-based methods cluster together with *1DCNN_E*, which has the most different representation out of them. In contrast, the models that do not use ELMo features learned very different embeddings, as their neighborhoods have nearly zero overlap both to each other and to the ELMo-based models.

**Fig. 3.**
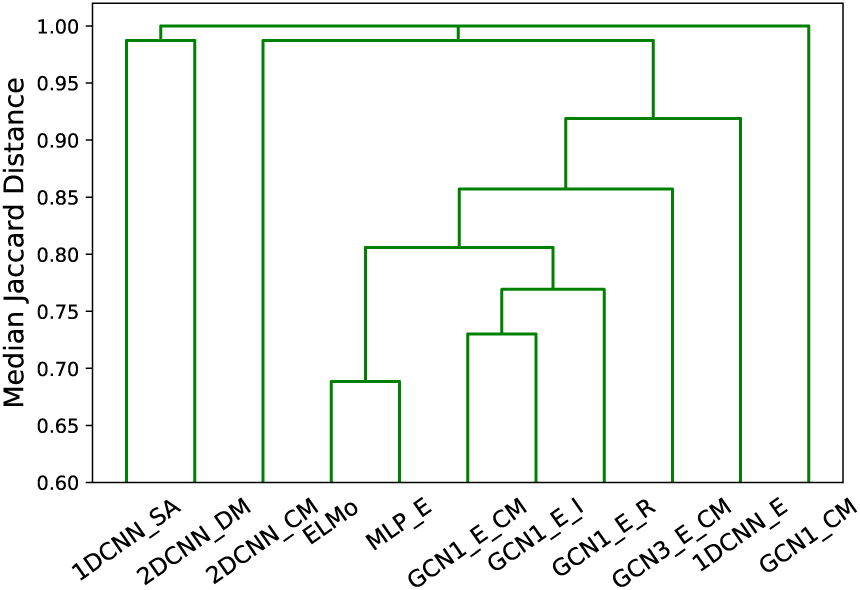
Hierarchical clustering of the models based on the similarity of the 40 nearest neighbors of each protein in the embedding space.

## 4 Discussion

Our work continues upon two recent studies involving protein representation learning (Heinzinger *et al*., 2019) and its combination with contact maps applied in AFP (Gligorijevic *et al*., 2019). We confirm the power of the unsupervised ELMo embeddings in capturing relevant biological information about proteins (Heinzinger *et al*., 2019). Simply embedding the proteins into the learned 1,024-dimensional space and applying the *k*-NN classifier led to better AFP performance than the two baseline methods (BLAST and naive), as well as several commonly used hand-crafted features such as one-hot encoding of amino acids, *k*-mer counts, secondary structure and backbone angles. This implies that the ELMo model was able to learn an embedding space in which the similarity between two proteins reflects functional similarity reasonably well, although it was only exposed to amino acid sequences and not to GO annotations. However, this representation only coarsely reflects protein function, as demonstrated by the poor performance of the *k*-NN classifier on the most specific terms.

As expected, we were able to improve the prediction accuracy achieved by the unsupervised embeddings by training supervised AFP methods on the embedding space. A set of logistic regression classifiers trained individually for each GO term achieved comparable S_*min*_ to the *k*-NN, while achieving significantly higher *ROCAUC* in the *PDB* dataset. Contrary to expectation, the GCN and CNN models trained on the amino acid-level embeddings extracted by ELMo were not able to outperform the logistic regression model in terms of *ROCAUC* and F*max*. They did outperform it in terms of S_*min*_, though (with the GCN being the best according to that metric), hinting that the logistic regression might be less effective for more specific GO terms. However, in the *SP* dataset, which is larger and contains more specific GO terms, the differences in S_*min*_ are less profound. Moreover, replacing the linear model (LR) with a non-linear one (MLP) gave a significant performance boost, considerably outperforming all others in *ROCAUC* and achieving state-of-the-art *CAFA* performance. Supervised training also resulted in a more consistent performance across all levels of GO term specificity. All in all, the competitive performance of the protein-level models highlights the power of the unsupervised ELMo embeddings.

In (Gligorijevic *et al*., 2019), the authors report on the superiority of a 3-layer GCN using amino acid embeddings from a pre-trained language model based on a LSTM network over BLAST and a 1D-CNN using a one-hot encoded amino acid representation. They attribute this superiority to the use of graph convolutions to model the protein 3D structure represented by contact maps. However, our experiments show that a 1D-CNN with strong amino acid embeddings is competitive with the GCN. Both convolutional models exhibited severe performance decline when replacing the ELMo embeddings with one-hot encoded amino acids. Based on these, we cannot exclude the possibility that the language model of (Gligorijevic *et al*., 2019) is by itself powerful enough to explain (most of) the increase in performance. If that is indeed the case, it would account for the fact that replacing the true contact map with a predicted one does not cause a significant drop in performance (Gligorijevic *et al*., 2019). To support this claim, we trained another GCN model from scratch, keeping the same architecture as our best GCN, but replacing the contact map by a graph with all nodes disconnected. The performance of that network was similar to that of the original (using the contact map). The same pattern was observed when replacing the contact map with a random graph (both at training and test time), clearly demonstrating that the contribution of the contact maps is rather small. This observation is interesting, as protein 3D structure is much more difficult and expensive to obtain than the sequence.

One of the hyperparameters of our networks was the number of fully-connected (FC) layers between the global pooling layer and the output FC layer for the classification. In our experiments, we tested our models with zero and one intermediate FC layer and used the validation *ROCAUC* to select the optimal for each model. In cases where the performance was similar, we chose to keep the simpler model for testing, as having fewer parameters makes it less prone to overfitting and more likely to better generalize on unseen proteins. A clear pattern emerged from this selection: for both GCN and 1D-CNN networks trained with ELMo embeddings, the extra FC layer was not required. On the other hand, for networks trained with one-hot encoded amino acid features or without any sequence features, the more complicated architecture was always selected. This means that in the feature space learned by the convolutional layers, the different classes (GO terms) are “more linearly separable” when ELMo embeddings are used and learning a simple mapping from that space to the output classes is enough for good performance. In the absence of “good” input features, it is harder for the convolutions to learn a “good” embedding space and as a result a more complex classifier is needed.

One can reasonably assume that also in the case of the one-hot features, it would be possible to learn a better (supervised) embedding space that only requires one linear classification layer. However, that would take a deeper architecture with more convolutional layers to enable us to discover more complicated patterns in protein sequences. This is problematic because the amount of available labeled data is not enough to train deep models with a larger number of parameters. Also, building a deeper model increases not only training time but also the man-hours spent deciding on the correct architecture and tuning the larger number of hyperparameters. To make matters worse, one would have to repeat almost the whole process from scratch if the task changes e.g. from function prediction to structure prediction. Unsupervised pre-training relieves part of that burden by creating only one complicated, deep sequence model to learn a meaningful feature representation of amino acids or proteins, which can then be fed to simpler classifiers to obtain competitive performance in several tasks without much effort (Heinzinger *et al*., 2019), as we demonstrated here.

Our experiments suggest that combining structure information in the form of a contact map with sequence information is not straightforward, especially when high-quality sequence features are available. Using a one-layer GCN did lead to a small improvement in S_*min*_, but at the cost of worse term-centric *ROCAUC* than the logistic regression baseline. Also, joining a one-dimensional and a two-dimensional CNN that independently extract sequence and contact map features, respectively, did not improve performance over the 1D-CNN applied to sequence data only. It is unlikely that contact maps do not contain any functional information, so our observations could have two possible explanations: either the ELMo embeddings contain three-dimensional structure information or we are still unable to leverage the full potential of contact maps.

To test the first hypothesis one could train a classifier that takes the amino acid-level features as inputs and predicts contacts between amino acid pairs. Such models already exist and do quite well in the CASP challenges by using physicochemical properties, the position-specific scoring matrix (PSSM) and predictions about secondary structure, solvent accessibility and backbone angles (Cheng and Baldi, 2007; Jones *et al*., 2015; Wang *et al*., 2017). By replacing these features with sequence embeddings as in (Bepler and Berger, 2019), we would expect a considerable improvement in the performance of these models.

On the contrary, finding a more effective way of using distance or contact maps is not trivial. Here, we considered a contact threshold of 10 Å by following previous studies (Gligorijevic *et al*., 2019), which is a more relaxed threshold than the one used in CASP challenges (8 Å), but also used alternative threshold strategies and obtained similar results. One could argue that the distance matrix is more informative and should be preferred, but our experiments did not confirm that. A different way of using distance maps in a GCN has been proposed by (Fout *et al*., 2017) to predict protein interfaces. Firstly, instead of using a fixed distance threshold, Fout et al. define each amino acid as being “in contact” with its *k* nearest residues, which creates a directed graph as the property of being someone’s nearest neighbor is not commutative. Moreover, the distances between the *k* nearest residues were smoothed with a Gaussian kernel and used as edge features over which a different set of filters was learned (Fout *et al*., 2017). Further research is required to resolve this issue.

In conclusion, this study shows that deep unsupervised pre-training of protein sequences is beneficial for predicting molecular function, as it can capture useful aspects of the amino acid sequences. We also showed that combining these sequential embeddings with contact map information does not yield significant performance improvements in the task, hinting that the embeddings may already contain 3D structural information.

## Acknowledgements

The authors would like to thank Dr. Elvin Isufi and Chirag Raman for their valuable comments and feedback.

