## Supplementary Material for "Unsupervised protein embeddings outperform hand-crafted sequence and structure features at predicting molecular function"

Amelia Villegas-Morcillo [*,1,†], Stavros Makrodimitris [*,2,3], Roeland C.H.J. van Ham [2,3], Angel M. Gomez [1], Victoria Sanchez [1] and Marcel J.T. Reinders [2,4]

[1]Dept. of Signal Theory, Telematics and Communications, University of Granada, Granada, Spain,
[2]Delft Bioinformatics Lab, Delft University of Technology, Delft, the Netherlands,
[3]Keygene N.V., Wageningen, the Netherlands and
[4]Leiden Computational Biology Center, Leiden University Medical Center, Leiden, the Netherlands.
[*]Equal contribution.
[†]To whom correspondence should be addressed.

#### Contents

**1 Protein representations** **2**

**2 Function prediction methods and training** **3**

**3 Alternative graph convolution operators** **7**

**4 *PDB* and *SP* datasets preprocessing** **9**

**5 Performance of all methods on the *PDB* dataset** **10**

**6 Performance of sequence-based models using the *SP* and *CAFA* datasets** **11**

**7 Term-centric performance and term specificity** **12**

**8 Bootstrapping results** **16**

**9 Principal Components Analysis of supervised embeddings** **21**

**10 Statistical significance of correlation between functional and embedding similarity** **22**

**11 Clustering based on embedding similarity** **24**

### 1 Protein representations

#### ELMo embeddings

The ELMo embeddings were extracted using a pre-trained unsupervised language model [1], based on the ELMo model [2]. It outputs amino acid-level features $\mathbf{X}^E \in \mathbb{R}^{L\times d}$ with $d$=1,024, which can be averaged into a protein-level representation $\mathbf{x}^E \in \mathbb{R}^d$ so that:

$$x_i^E = \frac{1}{L}\sum_{l=1}^{L} X_{l,i}^E \tag{1}$$

where $L$ is the protein sequence length and $i = 1, ..., d$ denotes each feature.

#### One-hot encodings

For the one-hot encoding representation, we also obtained amino acid-level features $\mathbf{X}^{1h} \in \{0,1\}^{L\times d}$ and protein-level features $\mathbf{x}^{1h} \in \mathbb{R}^d$ using the equation (1), which contain the frequency of each amino acid in the sequence. Here we considered $d$=26 features, i.e. the 20 amino acids 'ARNDC-QEGHILKMFPSTWYV' plus 6 special characters 'UOBZJX' defined by the FASTA format.

#### k-mer counts

We also used a protein-level representation based on $k$-mer counts, with $k$=[3,4,5]. As the dimensionality if this representation grows exponentially with $k$, we applied truncated Singular Value Decomposition (SVD), keeping the first $d = 1,024$ components in each case ($\mathbf{x}^{kmer} \in \mathbb{R}^d$). $k$=4 provided the best validation results.

#### Protein distance map

For each protein structure, we extracted the protein-level distance map by computing the Euclidean distances between all pairs of beta carbon atoms (alpha carbon atoms for Glycine). This real-valued matrix was then converted to fixed-range maps, more suitable for being processed by our neural network models. One is the binary contact map $\mathbf{A} \in \{0,1\}^{L\times L}$, obtained using a single threshold of 10 Å. We also considered a quantized distance map representation by using several thresholds/bins from 5 to 20 Å. We tried the following bins:

- 4 bins:
  - Distance thresholds: <5, [5-8), [8-15), [15-20), ≥20 Å
  - Bin values/weights: $[1, 0.75, 0.5, 0.25, 0]$
- 16 bins:
  - Distance thresholds: <5, [5-6), [6-7), ..., [18-19), [19-20), ≥20 Å
  - Bin values/weights: $[1, 15/16, 14/16, ..., 2/16, 1/16, 0]$
- 26 bins:
  - Distance thresholds: <5, [5-5.6), [5.6-6.2), ..., [18.8-19.4), [19.4-20), ≥20 Å
  - Bin values/weights: $[1, 25/26, 24/26, ..., 2/26, 1/26, 0]$

#### Structural features

We also tested an amino acid-level structural representation $\mathbf{X}^{SA} \in \mathbb{R}^{L\times d}$ with $d$=17. These features include the secondary structure (one-hot encoded 8-states 'HBEGITS-') and relative accessible surface area obtained from DSSP (Define Secondary Structure of Proteins) [3], along with the sine and cosine of the backbone angles $[\phi, \psi, \theta, \tau]$ [4].

### 2 Function prediction methods and training

#### Neural network architecture hyperparameters

In Table S1, the architectures of the MLP, 1D-CNN and 2D-CNN models are described. The architectures of the 3-layer and the 1-layer GCN's as well as that of the combination of the 1D-CNN and the 2D-CNN are shown on Table S2. Figures S1-S5 show a sketch of the architectures of 1D-CNN, 2D-CNN, 3-layer GCN, 1-layer GCN and 1D-CNN+2D-CNN respectively.

Table S1: Neural network hyperparameters for the MLP, 1D-CNN and 2D-CNN models. For each model, input feature information, architecture layers and names are provided. Parentheses indicate an optional intermediate FC layer followed by ReLU. The variable $C$ refers to the number of MFO GO terms to be classified by the network (which depends on the dataset).

| Model | MLP | 1D-CNN | 2D-CNN |
|---|---|---|---|
| Input | Protein-level features | Amino acid-level features | Contact map or<br>Quantized distance map |
| Layers | FC 512 - ReLU<br>FC $C$ - Sigmoid | 1Dconv 5x64 - ReLU<br>1Dconv 5x512 - ReLU<br>Global max pool 512<br>(FC 256 - ReLU)<br>FC $C$ - Sigmoid | 2Dconv 5x5x64 - ReLU<br>2Dconv 5x5x512 - ReLU<br>Global max pool 512<br>(FC 256 - ReLU)<br>FC $C$ - Sigmoid |
| Names | *MLP_{E,1h,kmer}* | *1DCNN_{E,1h,SA}* | *2DCNN_{CM,DM}* |

Table S2: Neural network hyperparameters for the 3-layer GCN, 1-layer GCN and the combined 1D-CNN with 2D-CNN models. For each model, input feature information, architecture layers and names are provided. Parentheses indicate an optional intermediate FC layer followed by ReLU. The symbol + indicates the combination of models. The variable $C$ refers to the number of MFO GO terms to be classified by the network (which depends on the dataset).

| Model | 3-layer GCN | 1-layer GCN | 1D-CNN + 2D-CNN |
|---|---|---|---|
| Input | Amino acid-level features and contact map (or quantized distance map) | | |
| Layers | Gconv 256 - ReLU<br>Gconv 256 - ReLU<br>Gconv 512 - ReLU<br>Global sum pool 512<br>(FC 256 - ReLU)<br>FC $C$ - Sigmoid | Gconv 512 - ReLU<br>Global sum pool 512<br>(FC 256 - ReLU)<br>FC $C$ - Sigmoid | 1Dconv 5x64 - ReLU<br>+ 2Dconv 5x5x64 - ReLU<br>1Dconv 5x512 - ReLU<br>+ 2Dconv 5x5x512 - ReLU<br>Global max pool 512 + 512<br>(FC 256 - ReLU)<br>FC $C$ - Sigmoid |
| Names | *GCN3_{E,1h,SA}_CM*<br>*GCN3_{E,1h}_DM* | *GCN1_{E,1h,SA}_CM*<br>*GCN1_{E,1h}_DM* | *1DCNN_1h+2DCNN_CM*<br>*1DCNN_E+2DCNN_CM* |

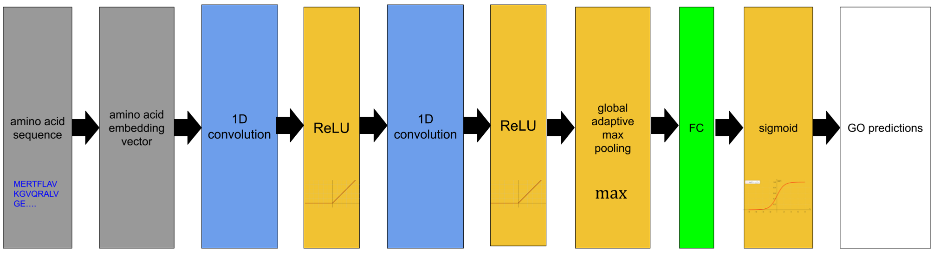


Figure S1: Neural network architecture for the 1D-CNN model.

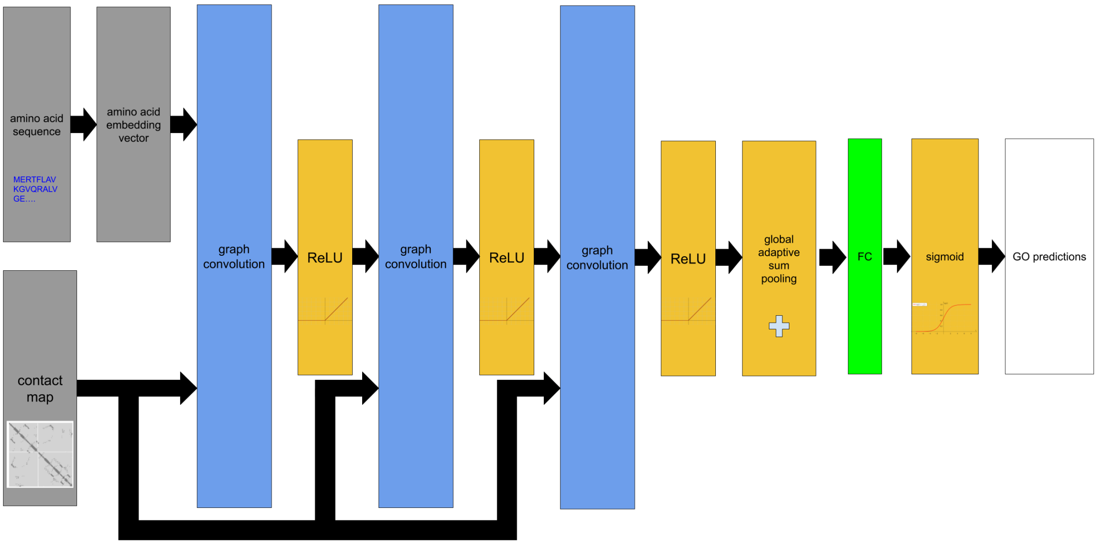


Figure S2: Neural network architecture for the 3-layer GCN model.

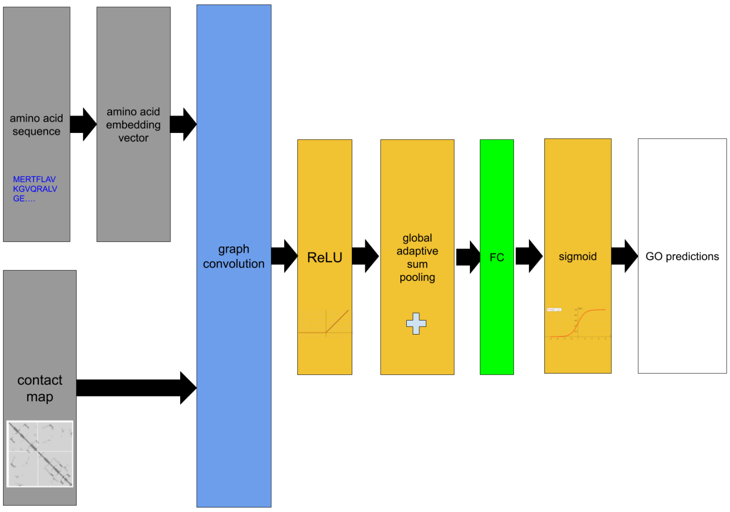


Figure S3: Neural network architecture for the 1-layer GCN model.

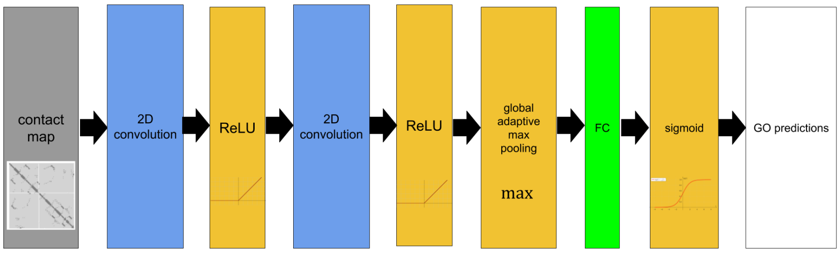


Figure S4: Neural network architecture for the 2D-CNN model.

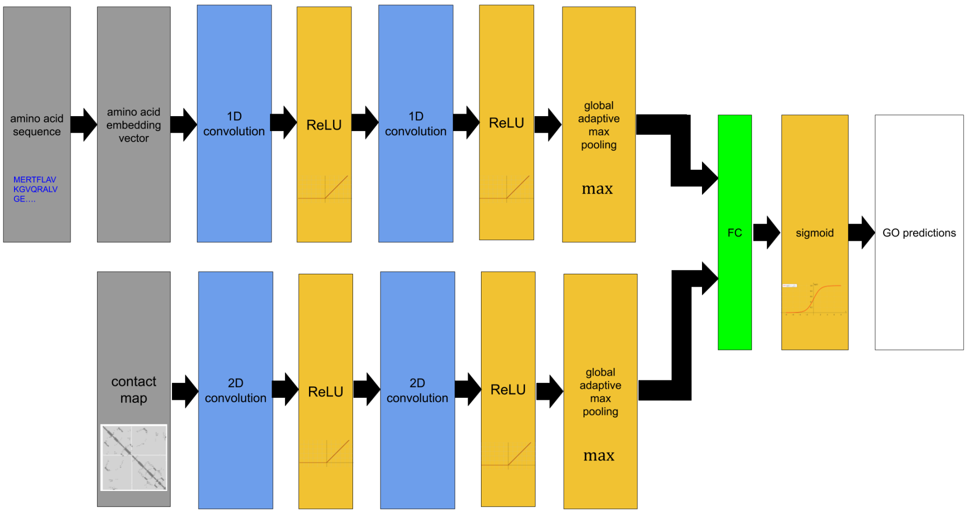


Figure S5: Neural network architecture for the combined 1D-CNN with 2D-CNN model.

#### Training details

For the $k$-NN classifier, we considered Euclidean distance and the number of nearest neighbors $k \in [1, 2, 3, 5, 7, 11, 15, 21, 25]$ was chosen using the validation set. For logistic regression, we trained an independent binary classifier for each GO term using L2 regularization. We used stochastic gradient descent (SGD) to accelerate the optimization. The optimal value for the penalty coefficient $\lambda$ was tuned jointly for all terms using the validation set. The values we tested were $10^{-3}$, $10^{-4}$ and $10^{-5}$.

The neural network models (MLP, 1D-CNN, 2D-CNN, GCN, and the combined 1D-CNN with 2D-CNN) were trained in a mini-batch mode for a variable number of epochs. We used a mini-batch size of 64, except for the 2D-CNN and the combined 1D-CNN with 2D-CNN models, in which we grouped protein samples of similar size together into mini-batches of sizes [1, 4, 8, 16, 32, 64] due to memory limitations. We trained all models by minimizing the average binary cross entropy over all GO terms. In order to prevent overfitting, we applied dropout [5] with drop probability 0.3 after the global pooling layer. For parameter updating we used the Adam optimizer [6] with an initial learning rate of $5 \cdot 10^{-4}$, which we reduced by a factor of 10 every time we observed that the validation loss did not improve for five consecutive epochs. We also used the validation *ROCAUC* to select the optimal set of parameters for each model. The neural network models were implemented in Pytorch [7] and Pytorch Geometric [8].

#### Validation results using the intermediate FC layer

We used the *PDB* validation set to decide if the second FC layer (with 512 units) is needed in each neural network model or not. Table S3 shows the *ROCAUC* values for the 1D-CNN, 3-layer GCN, 1-layer GCN and 1D-CNN + 2D-CNN models (considering one-hot encodings or ELMo embeddings), and the 2D-CNN model. The simpler model (with one FC layer) was selected if it was better or if the performance difference between the two options was smaller than 0.01.The simpler model architecture was preferred for all networks in the presence of ELMo embeddings and the more complex one when using one-hot encodings or contact map information only, with the exception of *1DCNN_1h*. This shows that functions are more linearly separable in the embedding spaces learned when using ELMo features.

Table S3: *ROCAUC* of neural network models trained with 1 and 2 FC layers on the *PDB* validation set. The results of the selected option for each model are shown in bold.

| Method | 1 FC layer | 2 FC layers |
|---|---|---|
| *1DCNN_E* | **0.929** | 0.924 |
| *1DCNN_1h* | **0.844** | 0.838 |
| *GCN3_E_CM* | **0.927** | 0.932 |
| *GCN3_1h_CM* | 0.826 | **0.873** |
| *GCN1_E_CM* | **0.924** | 0.930 |
| *GCN1_1h_CM* | 0.799 | **0.857** |
| *GCN1_CM* | 0.622 | **0.696** |
| *2DCNN_CM* | 0.795 | **0.847** |
| *1DCNN_E+2DCNN_CM* | **0.909** | 0.893 |
| *1DCNN_1h+2DCNN_CM* | **0.835** | 0.834 |

#### Validation results using quantized distance maps with different bins

We investigated alternative ways to process the distance map. To do so, we tested a quantized distance map representation using several thresholds/bins from 5 to 20Å. We used the *PDB* validation set to decide the optimal number of bins for each model. Table S4 shows the *ROCAUC* values for the 1-layer GCN (using ELMo embeddings, one-hot encodings or the degree as node features) and the 2D-CNN models, for 4, 16 or 26 bins. We kept the same network architecture as with contact map features (Table S3), i.e. using 2 FC layers in all models, except for the 1-layer GCN with ELMo embeddings (*GCN1_E_DM*). As shown in Table S4, the differences in the 1-layer GCN model are minimal, so we selected the 4 bins for the quantized distance map. For the 2D-CNN model, the 26-bin model clearly outperformed the others.

Table S4: *ROCAUC* of the 1-layer GCN and the 2D-CNN models trained using the quantized distance map with different number of bins (4, 16 or 26), on the *PDB* validation set. The results of the selected option for each model are show in bold.

| Method | *ROCAUC* 4 bins | *ROCAUC* 16 bins | *ROCAUC* 26 bins |
|---|---|---|---|
| *GCN1_E_DM* | **0.921** | 0.925 | 0.917 |
| *GCN1_1h_DM* | **0.861** | 0.866 | 0.866 |
| *GCN1_DM* | **0.669** | 0.694 | 0.680 |
| *2DCNN_DM* | 0.767 | 0.784 | **0.832** |

### 3 Alternative graph convolution operators

#### Definitions of operators

The graph convolution operator proposed in [9] is a polynomial approximation of the spectral graph convolution operation [10]. It is a first-order approximation, meaning that it makes use of Chebyshev polynomials up to degree one. That means that when calculating the new feature vector for the node using this approach, we only consider the current features of that node plus the features of its immediate neighbors (nodes that are 1 "hop" away). Higher-order approximations using Chebyshev polynomials were proposed in [11], leading to the Chebyshev spectral graph convolution operator defined as:

$$\mathbf{X}' = \sum_{k=0}^{K} \mathbf{Z}^{(k)}(\mathbf{X}, \hat{\mathbf{L}}) \cdot \mathbf{W}^{(k)} \tag{2}$$

In equation 2, $K$ is the order of the approximation and corresponds to the number of "hops" from the central node considered and $W^{(k)}$ is the weight matrix corresponding used to combine the features of nodes that are $k$ hops away from the central one. Finally, $\mathbf{Z}^{(k)}(\mathbf{X}, \hat{\mathbf{L}})$ is a Chebyshev polynomial of degree $k$ with $\mathbf{Z}^{(0)} = \mathbf{X}$, $\mathbf{Z}^{(1)} = \hat{\mathbf{L}} \cdot \mathbf{X}$ and $\mathbf{Z}^{(k)} = 2\hat{\mathbf{L}}\mathbf{Z}^{(k-1)} - \mathbf{Z}^{(k-2)}$ and $\hat{\mathbf{L}} = \frac{2}{\lambda_{max}}\mathbf{L} - \mathbf{I}$ is the normalized graph Laplacian.

The graph isomorphism network (GIN) is a form of spatial graph convolution [12]. If we use $(\mathbf{X})_i$ to denote the feature vector of node $i$, i.e. the $i$-th row of $\mathbf{X}$, the GIN uses the following operator:

$$(\mathbf{X}')_i = h\left((\mathbf{X})_i + \sum_{j \in N(i)} (\mathbf{X})_j, \mathbf{W}\right) \tag{3}$$

In equation 3, $h(\mathbf{x}, \mathbf{W})$ is a multi-layer perceptron with weights $\mathbf{W}$ and $N(i)$ is the set of nodes adjacent to node $i$.

The Gaussian mixture models (GMM) convolution operation uses a (hand-crafted) $d$-dimensional edge feature vector $\mathbf{u}(i, j) \in \mathbb{R}^d$ and defines a $J$-dimensional node similarity function $\mathbf{w}(\mathbf{u}) = (w_1(\mathbf{u}), w_2(\mathbf{u}), ..., w_J(\mathbf{u}))$ whose $m$-th element is calculated using equation 4 from the learnable parameters $\boldsymbol{\mu}_m$ and $\boldsymbol{\Sigma}_m$.

$$w_m(\mathbf{u}) = \exp\left(-\frac{1}{2}(\mathbf{u} - \boldsymbol{\mu}_m)^T \boldsymbol{\Sigma}_m^{-1}(\mathbf{u} - \boldsymbol{\mu}_m)\right) \tag{4}$$

#### Experiments and results

We tested the Chebyshev operator with $K = 2, 3, .., 10$ using one convolutional layer with 512 filters. In the validation set, we observed similar performance regardless of the value of $k$, so we only tested the models with $K = 2$ and $K = 10$. For the GIN model, we used a two-layer perceptron with 512 hidden units. For the GMM, we defined $u(i, j)$ as a single number denoting the linear distance in the sequence between residues $i$ and $j$, with $u(i, j) > 0$ if $i$ precedes $j$. The dimensionality of $w$ was also set to 1. To achieve the full potential of the GMM model, one needs to learn separate values for the mean and variance for every filter. However, due to memory limitations we were not able to train such a model and restricted ourselves to the simpler and less powerful version of the algorithm where $\boldsymbol{\mu}_m$ and $\boldsymbol{\Sigma}_m$ are shared across all filters.

We trained these models on the *PDB* dataset and tested on the corresponding test set. We compared different node features at the input: ELMo embeddings, one-hot encoding of the amino acids and no features (i.e. only information from the contact map). It should be noticed that we considered the model with an intermediate fully-connected layer when using either one-hot encodings or no features, while the simpler model was chosen when using ELMo embeddings. The results on the test set with at most 30% identity are listed in Table S5. We observed very similar performance by all models when using ELMo embeddings or one-hot encodings. However, the more complex graph convolutional operators (Chebyshev with $K = 10$ and GMM) perform better in terms of *ROCAUC* when processing only information from the contact map (without node features).

Table S5: Results of different graph convolutional operators on the *PDB* test subset with at most 30% sequence identity, using ELMo embeddings or one-hot encoded amino acids.

| Graph conv operator | ELMo embeddings | | One-hot encodings | |
|---|---|---|---|---|
| | $S_{min} \downarrow$ | $ROCAUC \uparrow$ | $S_{min} \downarrow$ | $ROCAUC \uparrow$ |
| Kipf GCN (1-layer) | 0.50 | 0.76 | 0.58 | 0.75 |
| Chebyshev ($K = 2$) | 0.51 | 0.75 | 0.57 | 0.75 |
| Chebyshev ($K = 10$) | 0.51 | 0.74 | 0.57 | 0.74 |
| GIN | 0.52 | 0.75 | 0.59 | 0.70 |
| GMM | 0.51 | 0.75 | 0.58 | 0.73 |

#### Conclusion

For contact map data combined with ELMo embeddings, all tested graph convolution operations perform similarly, so the inability to gain considerable improvement with respect to *1DCNN_E* is probably not due to choosing an unsuitable operation.

### 4 *PDB* and *SP* datasets preprocessing

#### Protein samples and GO terms

For the *PDB* dataset, we collected all Protein Data Bank (PDB) [13] identifiers with their corresponding annotations from the Gene Ontology (GO) knowledge base using the SIFTS resource (Structure Integration with Function, Taxonomy and Sequence) [14]. For *SP*, we used all amino acid sequences available in SwissProt in January 2020. In both cases, we considered proteins that were annotated with GO terms from the Molecular Function Ontology (MFO). We only used GO annotations derived by non-computational means, i.e. annotations with one of the following evidence codes: 'EXP', 'IDA', 'IPI', 'IMP', 'IGI', 'IEP', 'HTP', 'HDA', 'HMP', 'HGI', 'HEP', 'IBA', 'IBD', 'IKR', 'IRD', 'IC' and 'TAS'.

We retrieved protein structures from the RCSB PDB website [13] and obtained the individual chains with their corresponding amino acid sequences using PDB_Tool (`https://github.com/realbigws/PDB_Tool`) for the *PDB* dataset. Then, in both datasets, we only kept proteins with sequence length in the range [40, 1000]. The resulting *PDB* dataset contains in total 115,198 chains from 43,657 PDB proteins, while *SP* contains 90,227 protein sequences. We used CD-HIT [15] to group similar sequences, assigning proteins with more than 95% identity to the same cluster. From each cluster in the *PDB* dataset, we selected the chain with the highest-resolution structure, which could have been obtained with either X-ray crystallography, Nuclear Magnetic Resonance (NMR) or Cryo-Electron Microscopy. After this filtering step, we had a total of 11,749 sequences annotated with 3,918 MFO GO terms in *PDB*. We had 80,176 proteins with maximum 95% sequence identity in the *SP* dataset.

#### Training, validation and test sets

To ensure diversity in the evaluation, we imposed that all proteins in the *PDB* and *SP* test sets have at most 30% sequence identity to each other. For this purpose, we clustered all the protein sequences in each dataset with PSI-CD-HIT [15]. The clustering identified 2,971 sequences of maximally 30% identity in *PDB*, from which we randomly selected 1,175 to use as test set (10% of the entire dataset of 11,749 chains). The remaining data were randomly split into a training (80% of total chains) and a validation set (10% of total chains). The same approach was followed for *SP*, giving us 8,018 test proteins.

After this splitting, several GO terms ended up having too few or even no positive examples in at least one of the sets. We excluded from any further analysis all GO terms that had fewer than 40 positive examples in the training set or fewer than 5 in the validation or test sets. We then removed the proteins that were left with no annotations after this filtering. Finally, our training, validation and test sets for *PDB* had 9,395, 1,173 and 1,172 proteins respectively, annotated with $C$=256 MFO GO terms. In the *SP* dataset, we had 63,994 training, 8,004 validation and 8,001 test proteins, annotated with $C$=441 MFO GO terms. In order to assess the generalization ability of our trained models in evolutionarily distant proteins, we further defined a subset of the test set using BLAST, in which all proteins had sequence identity smaller than 30% to any of the training proteins. This test subset contains 450 protein chains for the *PDB* dataset and 3,530 sequences for the *SP*.

### 5 Performance of all methods on the *PDB* dataset

Table S6 shows the performance of the tested models on the *PDB* dataset based on $S_{min}$, $ROCAUC$ and $F_{max}$. We observed that models that use ELMo perform far better than their counterparts with one-hot encoded amino acids and that simple methods on the protein-level embeddings (*LR_E* and *MLP_E*) perform equally to models that use convolutions (*1DCNN_E*, *GCN3_E_CM* and *GCN1_E_CM*).

Table S6: $S_{min}$, $ROCAUC$ and $F_{max}$ of all methods on the 30% sequence identity *PDB* test set with 450 proteins and $C$=256 MFO GO terms. The values in brackets indicate 95% confidence intervals estimated using 1,000 bootstraps. The highest performance is indicated in boldface.

| Method | $S_{min}$ ↓ | $ROCAUC$ ↑ | $F_{max}$ ↑ |
|---|---|---|---|
| Naive | 0.61 [0.608, 0.620] | 0.50 [0.500, 0.500] | 0.43 [0.410, 0.451] |
| BLAST | 0.53 [0.512, 0.556] | 0.62 [0.597, 0.642] | 0.37 [0.346, 0.405] |
| *kNN_E* | 0.54 [0.519, 0.551] | 0.72 [0.701, 0.757] | 0.47 [0.451, 0.500] |
| *LR_E* | 0.55 [0.515, 0.549] | 0.77 [0.744, 0.796] | 0.50 [0.466, 0.512] |
| *MLP_E* | 0.51 [0.489, 0.524] | **0.82 [0.797, 0.839]** | **0.51 [0.483, 0.533]** |
| *1DCNN_E* | 0.51 [0.488, 0.525] | 0.77 [0.737, 0.807] | 0.50 [0.476, 0.526] |
| *GCN3_E_CM* | 0.52 [0.498, 0.534] | 0.74 [0.711, 0.776] | 0.48 [0.456, 0.503] |
| *GCN3_E_DM* | 0.52 [0.500, 0.536] | 0.77 [0.739, 0.797] | 0.47 [0.450, 0.499] |
| *GCN1_E_CM* | **0.50 [0.477, 0.515]** | 0.76 [0.724, 0.789] | **0.51 [0.492, 0.540]** |
| *GCN1_E_I* | 0.50 [0.483, 0.519] | 0.76 [0.723, 0.793] | — |
| *GCN1_E_R* | 0.51 [0.485, 0.524] | 0.77 [0.747, 0.798] | — |
| *GCN1_E_DM* | 0.51 [0.488, 0.526] | 0.73 [0.697, 0.765] | 0.49 [0.471, 0.524] |
| *kNN_1h* | 0.60 [0.588, 0.610] | 0.60 [0.572, 0.622] | 0.41 [0.394, 0.432] |
| *LR_1h* | 0.60 [0.592, 0.608] | 0.68 [0.641, 0.699] | 0.42 [0.401, 0.439] |
| *MLP_1h* | 0.59 [0.586, 0.602] | 0.71 [0.677, 0.735] | 0.43 [0.411, 0.450] |
| *1DCNN_1h* | 0.60 [0.593, 0.610] | 0.63 [0.587, 0.655] | 0.40 [0.382, 0.425] |
| *GCN3_1h_CM* | 0.58 [0.567, 0.591] | 0.71 [0.672, 0.734] | 0.41 [0.387, 0.429] |
| *GCN3_1h_DM* | 0.59 [0.570, 0.597] | 0.68 [0.651, 0.712] | 0.39 [0.370, 0.413] |
| *GCN1_1h_CM* | 0.58 [0.565, 0.588] | 0.75 [0.715, 0.780] | 0.43 [0.407, 0.449] |
| *GCN1_1h_I* | 0.59 [0.580, 0.600] | 0.65 [0.611, 0.683] | — |
| *GCN1_1h_R* | 0.59 [0.576, 0.595] | 0.70 [0.655, 0.727] | — |
| *GCN1_1h_DM* | 0.58 [0.571, 0.593] | 0.73 [0.699, 0.757] | 0.41 [0.388, 0.430] |
| *kNN_kmer* | 0.61 [0.599, 0.616] | 0.52 [0.496, 0.539] | 0.40 [0.381, 0.418] |
| *LR_kmer* | 0.65 [0.645, 0.652] | 0.66 [0.628, 0.684] | 0.41 [0.394, 0.429] |
| *MLP_kmer* | 0.61 [0.600, 0.613] | 0.63 [0.584, 0.658] | 0.35 [0.337, 0.371] |
| *1DCNN_SA* | 0.56 [0.542, 0.573] | 0.72 [0.680, 0.759] | 0.45 [0.427, 0.475] |
| *GCN3_SA_CM* | 0.57 [0.553, 0.580] | 0.67 [0.637, 0.716] | 0.42 [0.399, 0.441] |
| *GCN1_SA_CM* | 0.58 [0.560, 0.586] | 0.69 [0.658, 0.720] | 0.42 [0.406, 0.450] |
| *GCN1_CM* | 0.60 [0.589, 0.604] | 0.64 [0.605, 0.674] | 0.43 [0.412, 0.456] |
| *GCN1_DM* | 0.60 [0.589, 0.608] | 0.63 [0.594, 0.664] | 0.43 [0.412, 0.457] |
| *2DCNN_CM* | 0.58 [0.561, 0.592] | 0.68 [0.641, 0.709] | 0.41 [0.387, 0.433] |
| *2DCNN_DM* | 0.59 [0.568, 0.598] | 0.66 [0.623, 0.695] | 0.38 [0.362, 0.410] |
| *1DCNN_E+2DCNN_CM* | 0.54 [0.519, 0.553] | 0.74 [0.704, 0.775] | 0.46 [0.435, 0.487] |
| *1DCNN_1h+2DCNN_CM* | 0.60 [0.582, 0.605] | 0.61 [0.578, 0.648] | 0.39 [0.370, 0.412] |

### 6 Performance of sequence-based models using the *SP* and *CAFA* datasets

#### *SP* dataset

Table S7: $S_{min}$, $ROCAUC$ and $F_{max}$ of sequence methods on the *SP* test set with 3,530 proteins and $C$=441 MFO GO terms. The values in brackets indicate 95% confidence intervals estimated using 100 bootstraps. The best performance is indicated in boldface.

| Method | $S_{min}\downarrow$ | $ROCAUC\uparrow$ | $F_{max}\uparrow$ |
|---|---|---|---|
| Naive | 0.56 [0.548, 0.565] | 0.50 [0.500, 0.500] | 0.47 [0.463, 0.478] |
| BLAST | 0.52 [0.508, 0.530] | 0.67 [0.660, 0.679] | 0.39 [0.379, 0.407] |
| *kNN_E* | 0.46 [0.456, 0.472] | 0.76 [0.746, 0.769] | 0.52 [0.513, 0.531] |
| *LR_E* | 0.48 [0.469, 0.484] | 0.86 [0.849, 0.867] | 0.50 [0.488, 0.507] |
| *MLP_E* | **0.45 [0.435, 0.452]** | **0.87 [0.860, 0.876]** | **0.56 [0.547, 0.566]** |
| *1DCNN_E* | 0.46 [0.448, 0.464] | 0.84 [0.832, 0.849] | 0.54 [0.527, 0.546] |
| *kNN_1h* | 0.55 [0.547, 0.558] | 0.63 [0.616, 0.637] | 0.41 [0.399, 0.415] |
| *LR_1h* | 0.55 [0.549, 0.560] | 0.72 [0.713, 0.731] | 0.43 [0.425, 0.441] |
| *MLP_1h* | 0.55 [0.547, 0.558] | 0.75 [0.741, 0.763] | 0.43 [0.418, 0.434] |
| *1DCNN_1h* | 0.54 [0.531, 0.544] | 0.73 [0.720, 0.741] | 0.43 [0.424, 0.439] |

#### *CAFA* dataset

For the *CAFA* dataset, we used the CAFA assessment tool (`https://cafatools.github.io/`) provided by the organizers to ensure comparability with the methods participating in the challenge. This tool provides evaluation only on $F_{max}$. Table S8 lists the performance of our models as well as of the two baselines and the five best-performing models as reported in [16].

Table S8: $F_{max}$ of sequence methods on the *CAFA* test set with 454 proteins and $C$=679 MFO GO terms. The performance of BLAST and naive baselines is also given, along with that of the five highest scoring models. We did not implement or evaluate models with an asterisk, but copied the values published in [16].

| Method | $F_{max}\uparrow$ |
|---|---|
| Naive* | 0.33 |
| BLAST* | 0.42 |
| *kNN_E* | 0.50 |
| *LR_E* | 0.51 |
| *MLP_E* | 0.55 |
| *1DCNN_E* | 0.53 |
| CAFA3 rank 1* | 0.62 |
| CAFA3 rank 2* | 0.61 |
| CAFA3 rank 3* | 0.61 |
| CAFA3 rank 4* | 0.61 |
| CAFA3 rank 5* | 0.54 |

### 7 Term-centric performance and term specificity

We examined how the term-centric *ROCAUC* of different models varies according to term specificity using the 441 GO terms in the *SP* dataset. We used two metrics to quantify how specific a term is: 1) the maximum path length in the GO graph from that term to the ontology root and 2) the Resnik information content of the term, which is defined as $ic(g) = -log(\frac{1}{N}\sum_i^N I(y_i(g) = 1))$, i.e. the negative logarithm of a term's frequency in the training set.

Although *kNN_E* had better overall *ROCAUC* than *1DCNN_1h*, the top-performing model that uses one-hot encoded amino acids, Figures S6a-b show that this superiority is not evident for the more specific terms, as the *kNN_E* tends to perform badly for those terms (Figures S7a-b). However, the rest of the supervised models trained on ELMo embeddings (*LR_E*, *MLP_E* and *1DCNN_E*) do outperform *1DCNN_1h* for all levels of terms specificity (Figures S6c-h). These models demonstrate an increased performance for specific terms (Figures S7c-h).

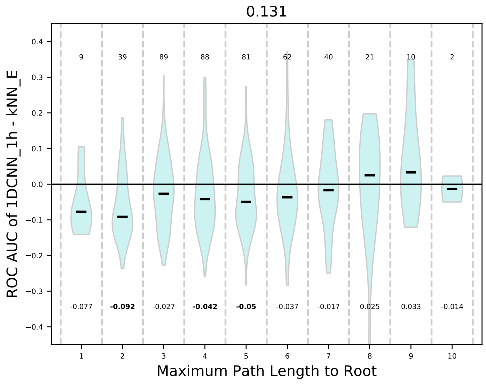


(a)

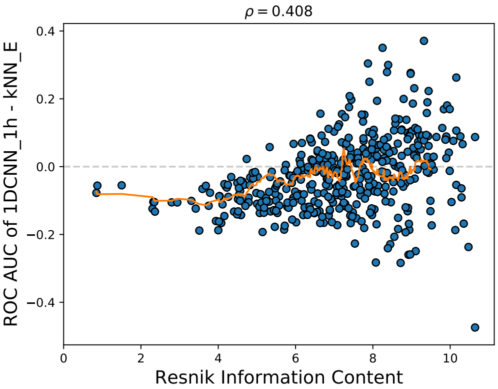


(b)

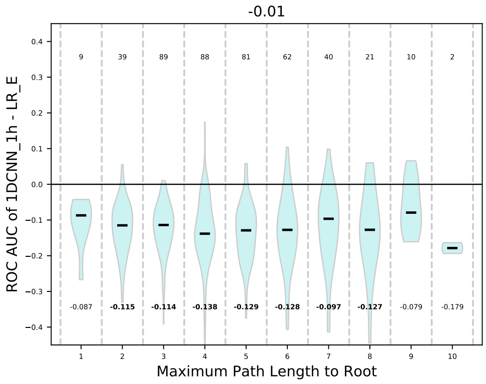


(c)

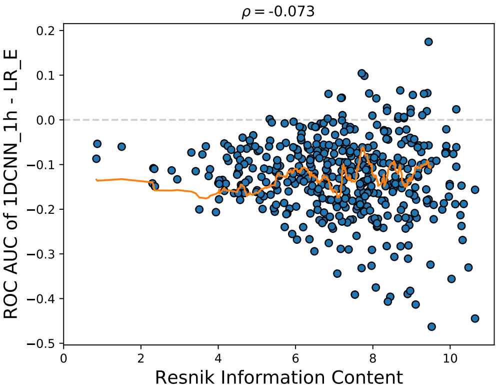


(d)

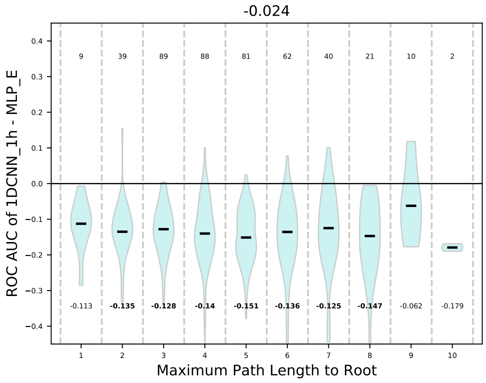


(e)

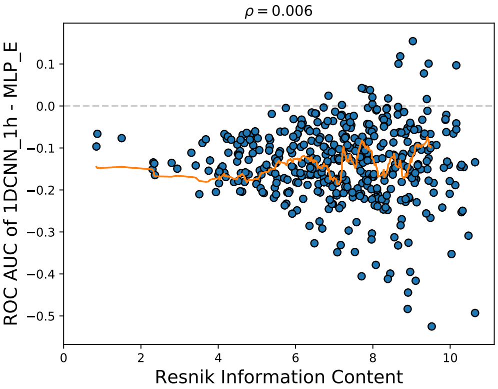


(f)

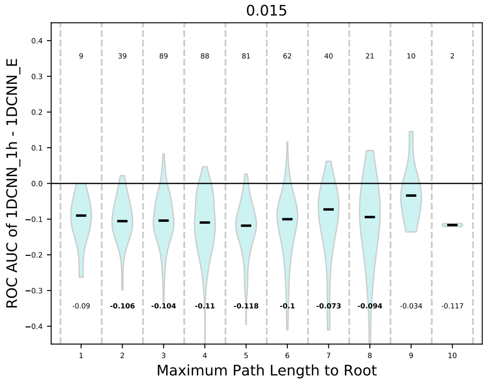


(g)

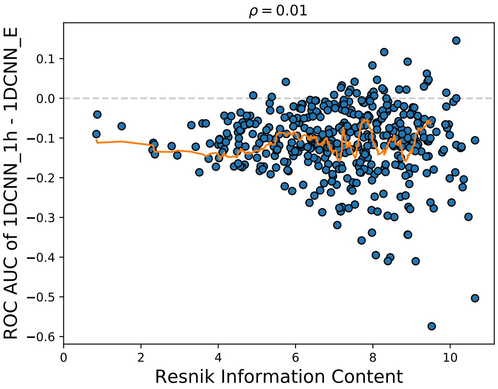


(h)

Figure S6: Difference in term-centric *ROCAUC* ($y-$ axis) between *1DCNN_1h* and *kNN_E* as a function of term specificity ($x-$axis). In (a), specificity is defined as the maximum path length to the ontology root. For each GO level we plot the distribution of the difference estimated with Gaussian kernels and show its median using a black dash. The value of the median is also shown as text below the plot, where bold values indicate that the difference is significant using a two-sided Wilcoxon rank sum test after Bonferroni multiple testing correction. The numbers above the distributions correspond to the number of GO terms at that specific level. In (b) specificity is defined as the Resnik information content. Every blue dot corresponds to a GO term and the orange line shows a local moving average calculated with a window of 30 terms. For both plots, the numbers at the top denote the Spearman rank correlation between the difference in *ROCAUC* and the term specificity and a line at $y = 0$ denotes equal performance for both models.

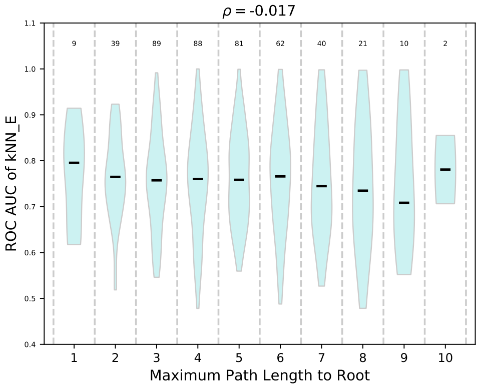
ρ = -0.017
ROC AUC of kNN_E
Maximum Path Length to Root
9 39 89 88 81 62 40 21 10 2

(a)

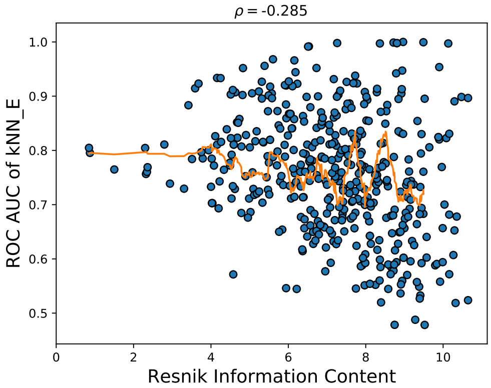
ρ = -0.285
ROC AUC of kNN_E
Resnik Information Content

(b)

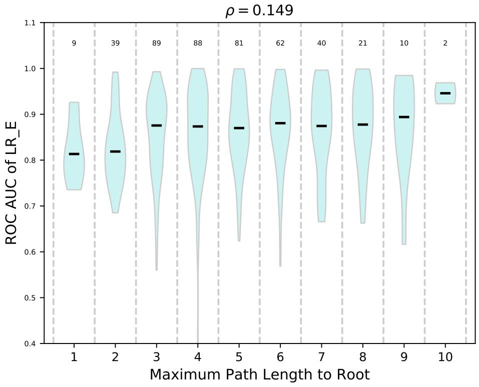
ρ = 0.149
ROC AUC of LR_E
Maximum Path Length to Root
9 39 89 88 81 62 40 21 10 2

(c)

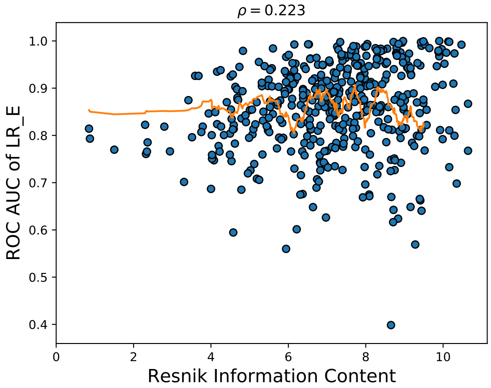
ρ = 0.223
ROC AUC of LR_E
Resnik Information Content

(d)

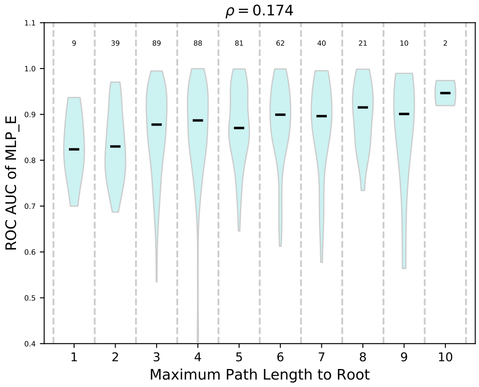
ρ = 0.174
ROC AUC of MLP_E
Maximum Path Length to Root
9 39 89 88 81 62 40 21 10 2

(e)

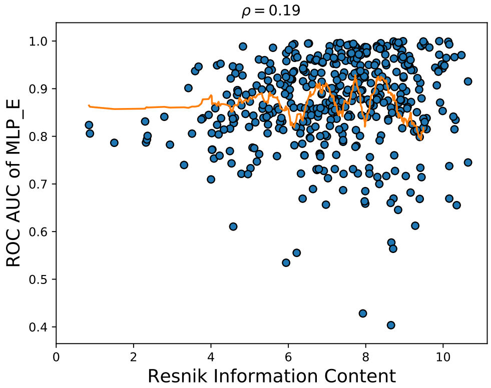
ρ = 0.19
ROC AUC of MLP_E
Resnik Information Content

(f)

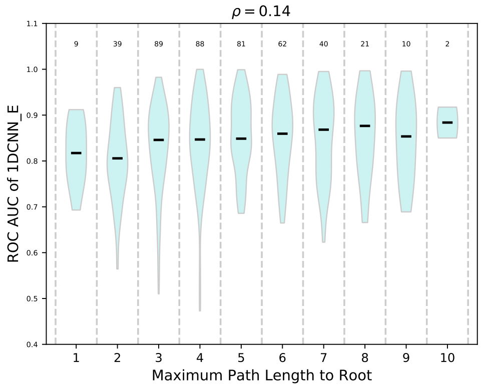


(g)

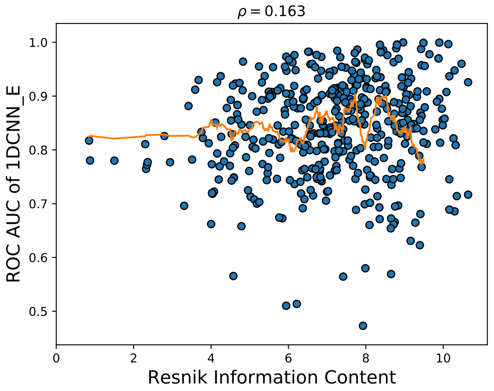


(h)

Figure S7: Difference in term-centric $ROCAUC$ ($y-$ axis) between *1DCNN_1h* and *kNN_E* as a function of term specificity ($x-$axis). In (a), specificity is defined as the maximum path length to the ontology root. For each GO level we plot the distribution of the difference estimated with Gaussian kernels and show its median using a black dash. The value of the median is also shown as text below the plot, where bold values indicate that the difference is significant using a two-sided Wilcoxon rank sum test after Bonferroni multiple testing correction. The numbers above the distributions correspond to the number of GO terms at that specific level. In (b) specificity is defined as the Resnik information content. Every blue dot corresponds to a GO term and the orange line shows a local moving average calculated with a window of 30 terms. For both plots, the numbers at the top denote the Spearman rank correlation between the difference in $ROCAUC$ and the term specificity and a line at $y = 0$ denotes equal performance for both models.

### 8 Bootstrapping results

#### Individual methods

We performed 1,000 bootstraps to estimate the variance of the observed performance values in the *PDB* dataset, by resampling proteins with replacement from the test set 1,000 times to create 1,000 bootstrap test sets and then evaluated each model in all test sets. Figure S8 shows the bootstrap results for $S_{min}$ and $ROCAUC$ for all methods.

#### Pairwise comparisons of methods

We also used these bootstraps to compare two methods with each other, by plotting the distribution of the performance difference between two methods over the 1,000 bootstrap test sets. These results for all pairs of models are shown in Figures S9-S11.

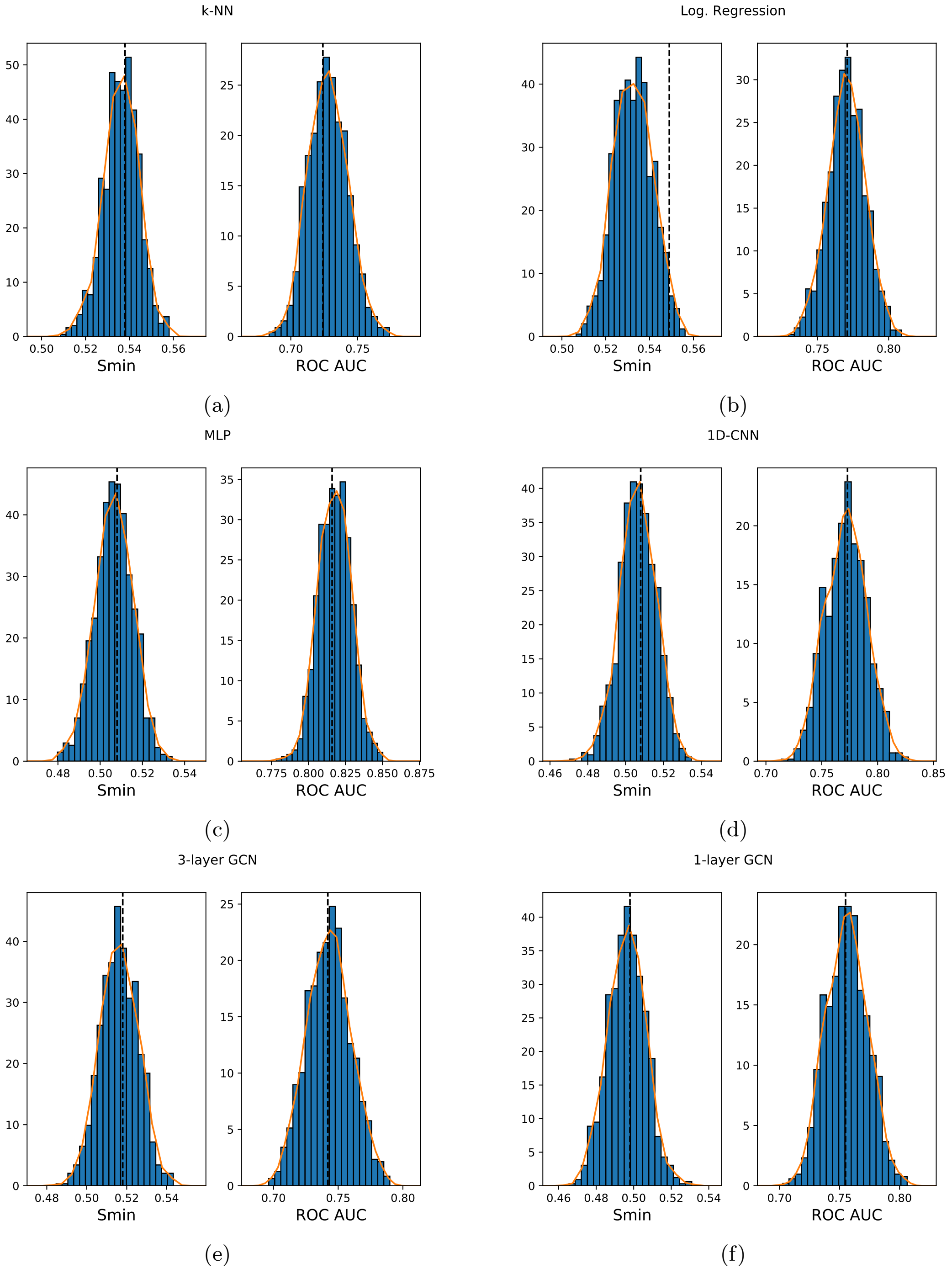


Figure S8: Distribution of the expected $S_{min}$ and $ROCAUC$ of different models in future test sets from the same distribution estimated using 1,000 bootstraps. The models are (a) *kNN_E*, (b) *LR_E*, (c) *MLP_E*, (d) *1DCNN_E*, (e) *GCN3_E_CM*, and (f) *GCN1_E_CM*. The distribution is shown both as a histogram (blue bars) and a density (orange curves), estimated with Gaussian kernels. A dashed vertical line denotes the model's performance on the actual test set.

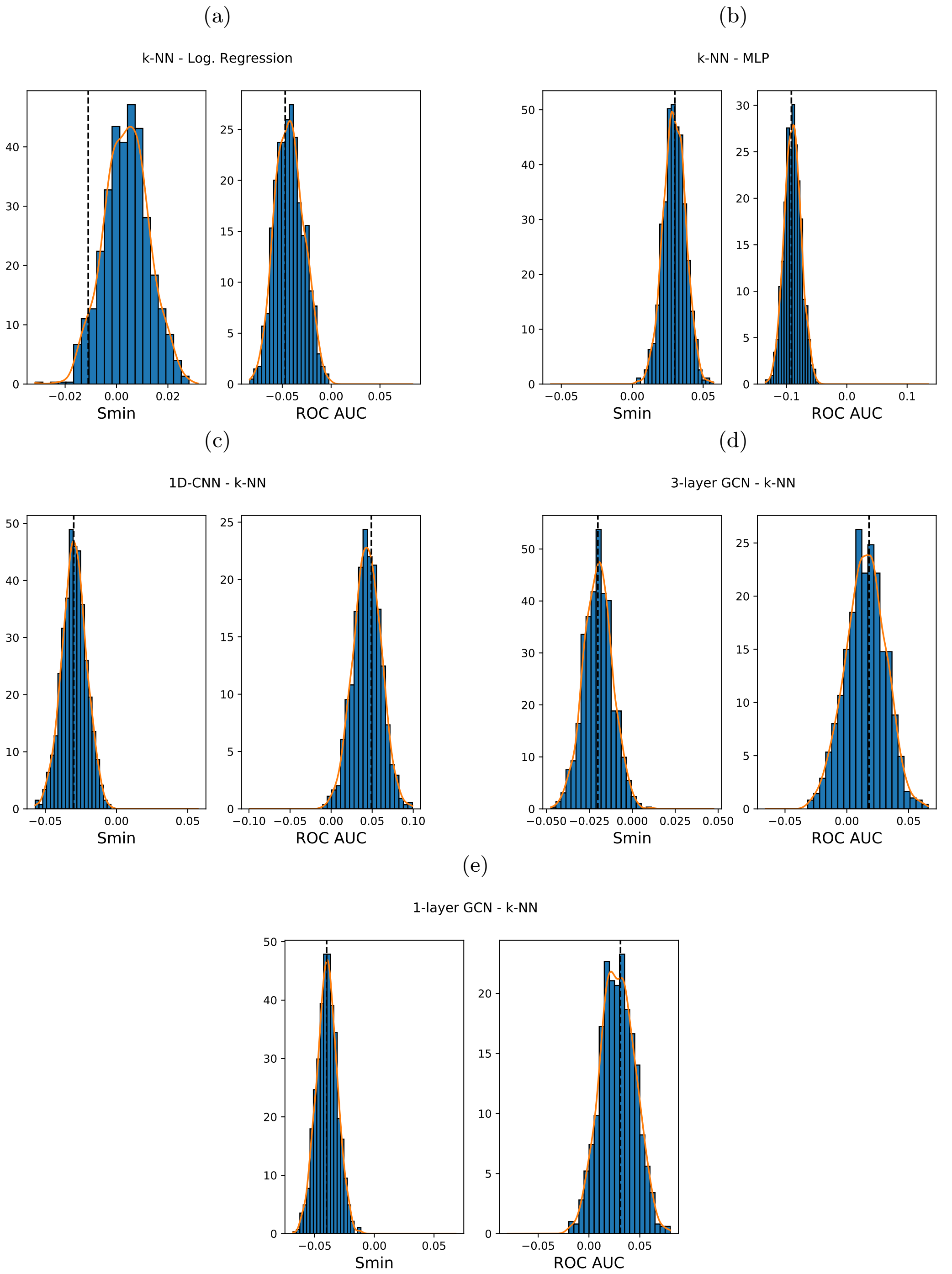


Figure S9: Distribution of the expected difference in $S_{min}$ and $ROC AUC$ between pairs of models in future test sets from the same distribution estimated using 1,000 bootstraps. The compared models are (a) *k-NN_E – LR_E*, (b) *kNN_E – MLP_E*, (c) *1DCNN_E – kNN_E*, (d) *GCN3_E_CM – kNN_E*, and (e) *GCN1_E_CM – kNN_E*. The distribution is shown both as a histogram (blue bars) and a density (orange curves), estimated with Gaussian kernels. A dashed vertical line denotes the performance difference on the actual test set.

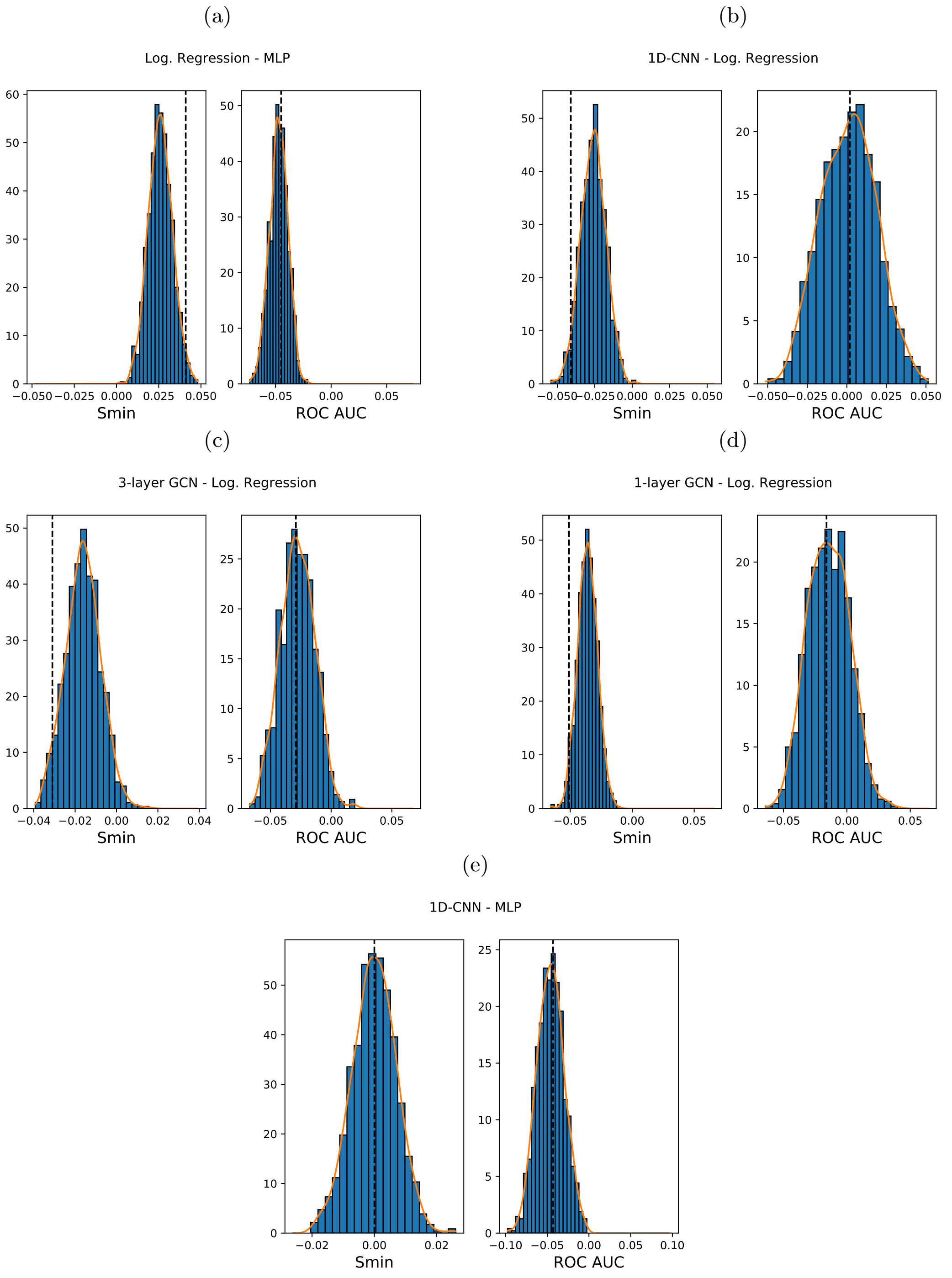


Figure S10: Distribution of the expected difference in $S_{min}$ and $ROCAUC$ between pairs of models in future test sets from the same distribution estimated using 1,000 bootstraps. The compared models are (a) *LR_E – MLP_E*, (b) *1DCNN_E – LR_E*, (c) *GCN3_E_CM – LR_E*, (d) *GCN1_E_CM – LR_E*, and (e) *1DCNN_E – MLP_E*. The distribution is shown both as a histogram (blue bars) and a density (orange curves), estimated with Gaussian kernels. A dashed vertical line denotes the performance difference on the actual test set.

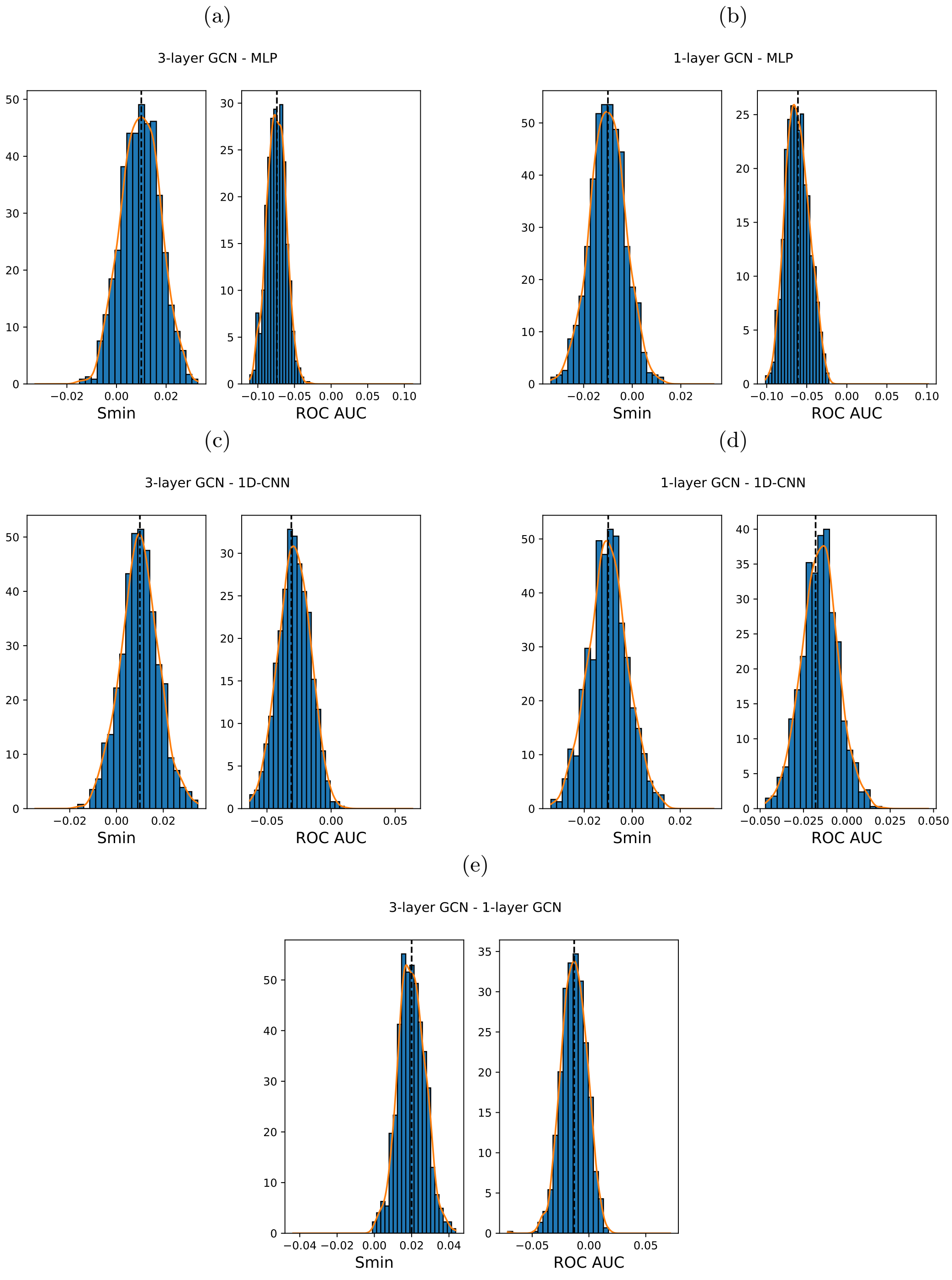


Figure S11: Distribution of the expected difference in $S_{min}$ and $ROCAUC$ between pairs of models in future test sets from the same distribution estimated using 1,000 bootstraps. The compared models are (a) *GCN3_E_CM – MLP_E*, (b) *GCN1_E_CM – MLP_E*, (c) *GCN3_E_CM – 1DCNN_E*, (d) *GCN1_E_CM – 1DCNN_E*, and (e) *GCN3_E_CM – GCN1_E_CM*. The distribution is shown both as a histogram (blue bars) and a density (orange curves), estimated with Gaussian kernels. A dashed vertical line denotes the performance difference on the actual test set.

### 9 Principal Components Analysis of supervised embeddings

We compared the supervised embeddings learned by the *MLP_E*, *1DCNN_E*, *1DCNN_SA*, *GCN3_E_CM*, *GCN1_E_CM*, *GCN1_CM*, *2DCNN_CM* and *2DCNN_DM* methods. We applied principal components analysis to the matrix obtained by inputting all available proteins in the *PDB* dataset to each network and recording the 512-dimensional feature vector before the last fully-connected layer. The rank of these matrices, which corresponds to the number of linearly independent features, is shown in Table S9.

Table S9: Rank of the embedding matrices obtained for all available proteins in the *PDB* dataset. The size of the learned embeddings by all methods is 512.

| Method | Embedding rank |
|---|---|
| *MLP_E* | 508 |
| *1DCNN_E* | 511 |
| *1DCNN_SA* | 512 |
| *GCN3_E_CM* | 512 |
| *GCN1_E_CM* | 512 |
| *GCN1_CM* | 105 |
| *2DCNN_CM* | 310 |
| *2DCNN_DM* | 512 |

The rank of the embedding matrix also corresponds to the number of principal components with non-zero eigenvalues, i.e. the components that explain part of the variance of the data. Figure S12 shows the relative variance explained by using more principal components. It is evident that the line for *GCN1_CM* starts at about 89%, which implies that the first principal component already explains most of the observed variance. The other models learn a lot more diverse representations (Figure S12), the most diverse being the *1DCNN*'s, *GCN1_E_CM* and *2DCNN_DM*, which require over 300 principal dimensions to explain 90% of the variance and the least being *MLP_E*, which requires fewer than 150.

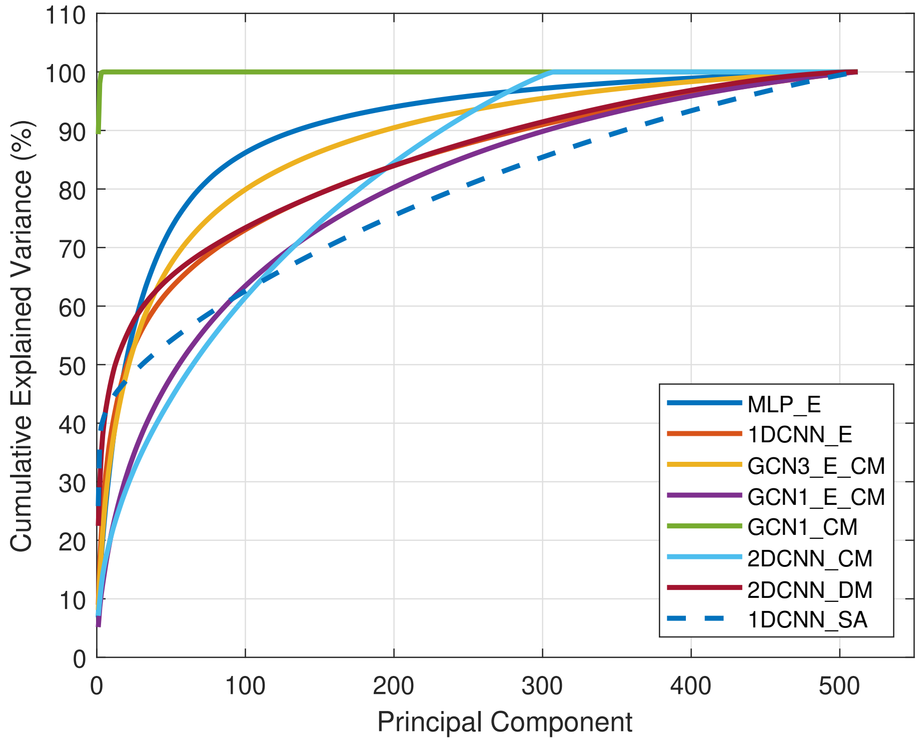


Figure S12: Cumulative percentage of explained variance ($y-$axis) as a function of the number principal components used ($x-$axis) of the supervised embeddings of different models. Each model is denoted by a different color.

#### 10 Statistical significance of correlation between functional and embedding similarity

We calculated rank correlations between embedding similarities and functional Jaccard similarities of proteins [17]. Because we are using all pairwise combinations of training and test proteins, the different observations of similarity are not independent, which means that our data do not meet the assumptions for calculating a p-value for the observed correlation. Therefore, we used permutation tests to assess significance.

##### Testing if observed correlation is significantly larger than zero

For this test, we randomly permuted the GO annotations of the test proteins, by shuffling the rows of the corresponding label matrix. That way, the functional similarity between a training and a test protein is randomized and no longer reflects their functions. We repeated this process 10,000 times, each time calculating the Spearman correlation between embedding similarity and randomized functional similarity. We thus obtained 10,000 samples from the distribution of the correlation under the null hypothesis, which we then compared to our observed value of $\rho = 0.07$. Figure S13 shows the null distribution and the observed correlation. In all 10,000 permutations, the null correlation was smaller than the observed one, meaning that the two similarities are significantly correlated with p-value $< 10^{-4}$.

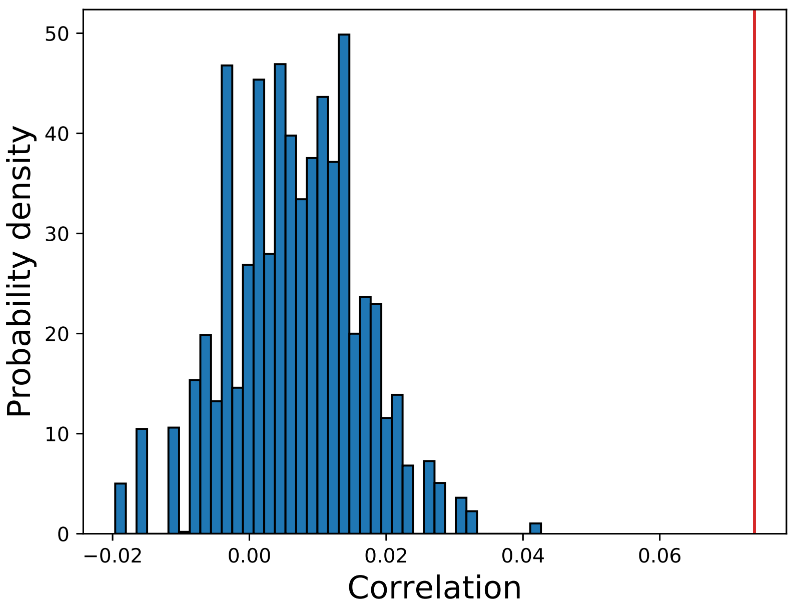


Figure S13: Distribution of Spearman correlations between functional Jaccard similarity and ELMo embedding similarity under the null hypothesis that the two similarities are uncorrelated. The null distribution was estimated using a permutation approach with 10,000 permutations. The red vertical line denotes the truly observed correlation value.

##### Testing if difference between two observed correlations is significantly larger than zero

To test if correlation values observed for two different embedding similarities (e.g. ELMo and *1DCNN_E*) are significantly different from each other, we performed a different test. The GO annotations of proteins and their pairwise semantic similarities were kept constant. Instead, we randomly exchanged the embedding similarity values for a subset of protein pairs, calculated the two new correlations and kept their signed difference. We repeated this 10,000 times to estimate the null distribution of the correlation difference and compared it to the observed correlation difference.

Figure S14 shows the null distributions and the observed differences in correlation values. We found that embedding similarity based on either *1DCNN_E*, *MLP_E* or *GCN1_E_CM* was significantly more correlated to functional similarity than the embedding similarity based on the unsupervised ELMo embeddings (permutation p-value $< 10^{-4}$ for all three cases).

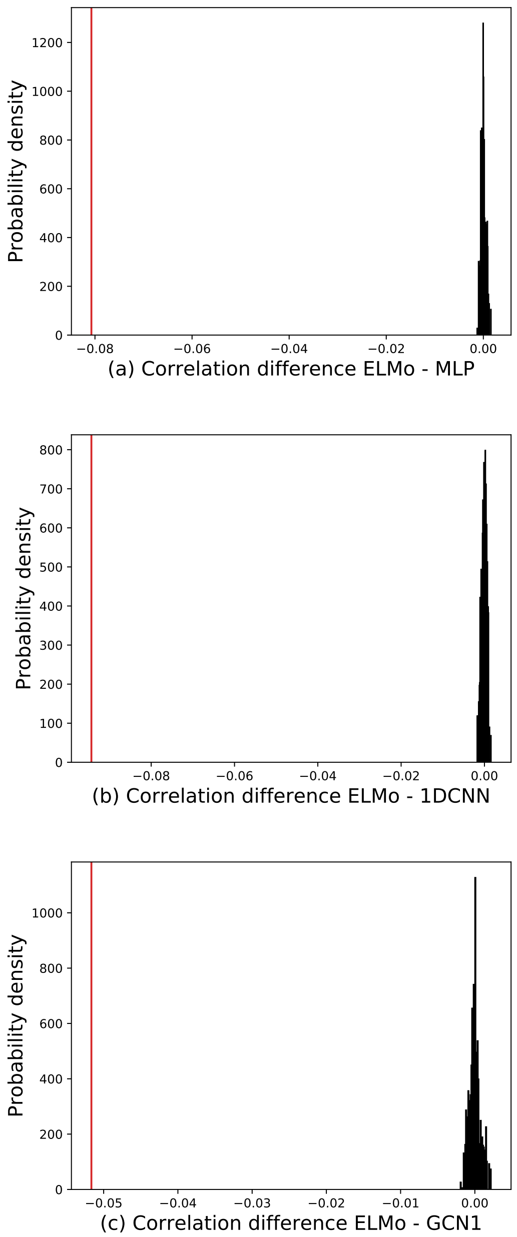


Figure S14: Distribution of the difference in Spearman correlations between functional Jaccard similarity and embedding similarity for a pair of embedding similarities under the null hypothesis that the difference in correlation is zero. The null distribution was estimated using a permutation approach with 10,000 permutations. The red vertical line denotes the truly observed difference in correlation values. In (a), the difference between ELMo and *MLP_E*, in (b) ELMo and *1DCNN_E* and in (c) ELMo and *GCN1_E_CM*.

#### Conclusion

These tests show that the unsupervised ELMo embeddings have already "learned" aspects of protein function and that further supervised training gives significantly more information.

#### 11 Clustering based on embedding similarity

For every *PDB* test protein, we found its 40 nearest training proteins in the embedding space using cosine similarity. Then we computed the Jaccard distance between the neighborhoods found for each test protein using two different embeddings. This gave us a distribution of neighborhood dissimilarities for each pair of embedding types. We used the median of this distribution as a measure of distance between embeddings and applied hierarchical clustering with complete linkage to group similar embeddings together.

[11] Michaël Defferrard, Xavier Bresson, and Pierre Vandergheynst. "Convolutional neural networks on graphs with fast localized spectral filtering". In: *Advances in Neural Information Processing Systems*. 2016. arXiv: 1606.09375.
